# Nuclear actin and DNA replication stress regulate the recruitment of human telomerase to telomeres

**DOI:** 10.1101/2024.03.25.586711

**Authors:** Ashley Harman, Melissa Kartawinata, Nohad M. Maroun, Darren R. Nguyen, William E. Hughes, Kevin Winardi, Anthony J. Cesare, Noa Lamm, Tracy M. Bryan

## Abstract

The recruitment of telomerase to telomeres is a tightly regulated process which is stimulated by replication stress and mediated by the DNA damage response regulatory kinase ATR. Here, we demonstrate that nuclear filamentous actin is important for telomerase recruitment under endogenous and replication stress conditions in immortal human cells. Inhibition of nuclear actin polymerization decreases the presence of telomerase at telomeres. This process is regulated by both ATR and mTOR kinases, and employs other regulators of actin structure and function, such as WASP, ARP2/3 and myosin. Nuclear filamentous actin serves as a site for telomerase recruitment, which is mediated by telomere tethering on actin fibres in response to replication stress, allowing telomerase to localize to telomeres containing stalled replication forks. Overall, these data demonstrate that, in human cells which express telomerase, telomeric replication stress triggers the recruitment of telomerase to telomeres via a nuclear actin network, enabling telomere length maintenance.

## INTRODUCTION

Telomeres are nucleoprotein complexes comprised of tandem DNA repeats, TTAGGG in vertebrates, which cap the ends of linear chromosomes to maintain the integrity of the genome and ensure cellular survival^1^. Given their resemblance to damaged DNA, exposed telomeres activate the DNA damage response (DDR) which can result in chromosome end-to-end fusions. To supress this response, telomeres are bound by the six-protein complex shelterin, which includes TRF1 and TRF2, which directly bind to duplex telomeric repeats and mediate binding of the remaining subunits: POT1, TPP1, RAP1 and TIN2^2^. Shelterin proteins perform various functions to maintain telomere structure and function, including regulating telomere length.

In human stem and germline cells and ∼85% of cancers, *de novo* telomere synthesis is performed by the ribonucleoprotein enzyme telomerase^3^. Human telomerase associates with a large number of different proteins^4^; however, the minimal catalytic core is the telomerase reverse transcriptase (hTERT), telomerase RNA (hTR) and the accessory protein dyskerin^5^. Recruitment of telomerase to telomeres is a highly regulated process, predominantly occurring during S phase^6,7^, which is at least partially regulated by TRF1^8^. TPP1 is also critical for telomerase recruitment, as it binds hTERT via an exposed N-terminal domain termed the TEL patch^9^. The interactions between TPP1, POT1 and TIN2 are also necessary to facilitate telomerase recruitment and processivity at telomeres^10–12^.

If telomeres become critically short, they can trigger activation of a DDR through either ATR or ATM^13^, PIKK (phosphatidylinositol-3 kinase-related) family kinases which govern the DDR. Shelterin typically represses the activation of both ATR and ATM at telomeres, specifically via POT1 and TRF2, respectively^14–17^. Furthermore, replication fork stalling within the telomere results in ATR activation^18^. Almost paradoxical, however, is the emerging evidence that the DDR is also crucial for regulation of telomerase presence at telomeres. In yeast, telomerase preferentially extends the shortest telomeres in a manner dependent on Tel1 (the yeast homolog of ATM)^19–22^. Furthermore, Tel1 is required for telomerase recruitment in budding yeast^23^, and both Tel1 and Rad3 (the fission yeast homolog of ATR) are required in fission yeast^24–26^. In human cells, ATR or ATM deficiency or mutation results in telomere shortening or instability^27,28^. This may rely upon TRF1 displacement, as phosphorylation of TRF1 at an ATM/ATR target site results in its depletion from telomeres and heightened telomerase recruitment^8,29^. Furthermore, the depletion or inhibition of ATR or ATM results in reduced telomerase presence at telomeres^8,30^. This axis is supported by the finding that induced replication stress promotes telomerase recruitment and ultimately causes telomere lengthening^8,18,31^. These studies together suggest a tightly regulated process for telomerase recruitment which is coordinated by DNA replication, the DDR and the telomere maintenance machinery.

An emerging body of research has illustrated that a key regulator of the DDR is nuclear filamentous actin (F-actin). Although classically considered cytoplasmic, nuclear F-actin forms in response to double strand breaks and replication stress, where it facilitates the DDR by re-localization of damaged sites to the nuclear periphery for repair or fork restart^32–37^. This occurs globally in response to DNA damage, but also specifically at telomeres undergoing replication stress^34,38^. Under replication stress conditions, the polymerization of nuclear F-actin is dependent on ATR, whose activity is required for downstream phosphorylation of another PIKK family kinase, mTOR^34^, in turn regulating F-actin through the Wiskott-Aldrich syndrome protein (WASP) family^39^.

The convergence of these data suggests there is controlled actin-mediated interplay between DNA replication, the DDR, and telomere maintenance. Here we demonstrate that nuclear F-actin regulates telomerase recruitment in an ATR- and mTOR-dependent manner in response to replication stress. Specifically, nuclear actin filaments appear to act as sites for telomerase recruitment, potentially due to telomere tethering on F-actin under conditions of replication stress. Telomeres are also re-localized toward the nuclear periphery via F-actin in response to replication stress. Furthermore, the recruitment of telomerase to telomeres occurs in proximity to stalled replication forks and F-actin.

## RESULTS

### F-actin polymerization facilitates telomerase recruitment to telomeres

To better visualize hTR presence at telomeres, we modified an existing fluorescence *in situ* hybridization (FISH) protocol^8,40^ by generating SABER (<u>s</u>ignal <u>a</u>mplification <u>b</u>y <u>e</u>xchange reaction) probes^41^ targeting both hTR and telomere sequences, greatly amplifying the intensity of the foci (Figures S1A-B). Cells were also labelled with EdU to identify S phase cells for quantification, as this is predominantly when telomerase recruitment to telomeres occurs within the cell cycle^6–8^.

Given that the recruitment of mammalian telomerase to telomeres is promoted by replication stress^8,42^, and the emerging understanding that F-actin is involved in resolution of stalled replication forks^34,37,43^, we explored whether actin polymerization is also involved in telomerase recruitment. Treatment of HEK293T cells with two inhibitors of actin polymerization, latrunculin A and latrunculin B (LatA and LatB)^44,45^, resulted in a significant reduction in telomerase recruitment to telomeres in S phase cells (Figures 1A-1C). Furthermore, while short term treatment (30 min) with the polymerase inhibitor aphidicolin (APH) is sufficient to induce a DNA damage response (as measured by phosphorylation of the ATR substrate Chk1; Figure 1D) and promote an increase in telomerase recruitment, this increased recruitment was prevented by inhibiting actin polymerization using either LatA or LatB (Figure 1B). Treatment with the latrunculins (with or without APH) did not cause substantial perturbation to cell cycle progression (Figure S1C), suggesting that the observed effects on telomerase are due to F-actin facilitating recruitment, and not due to fluctuations in the cell cycle.

**Figure 1.**
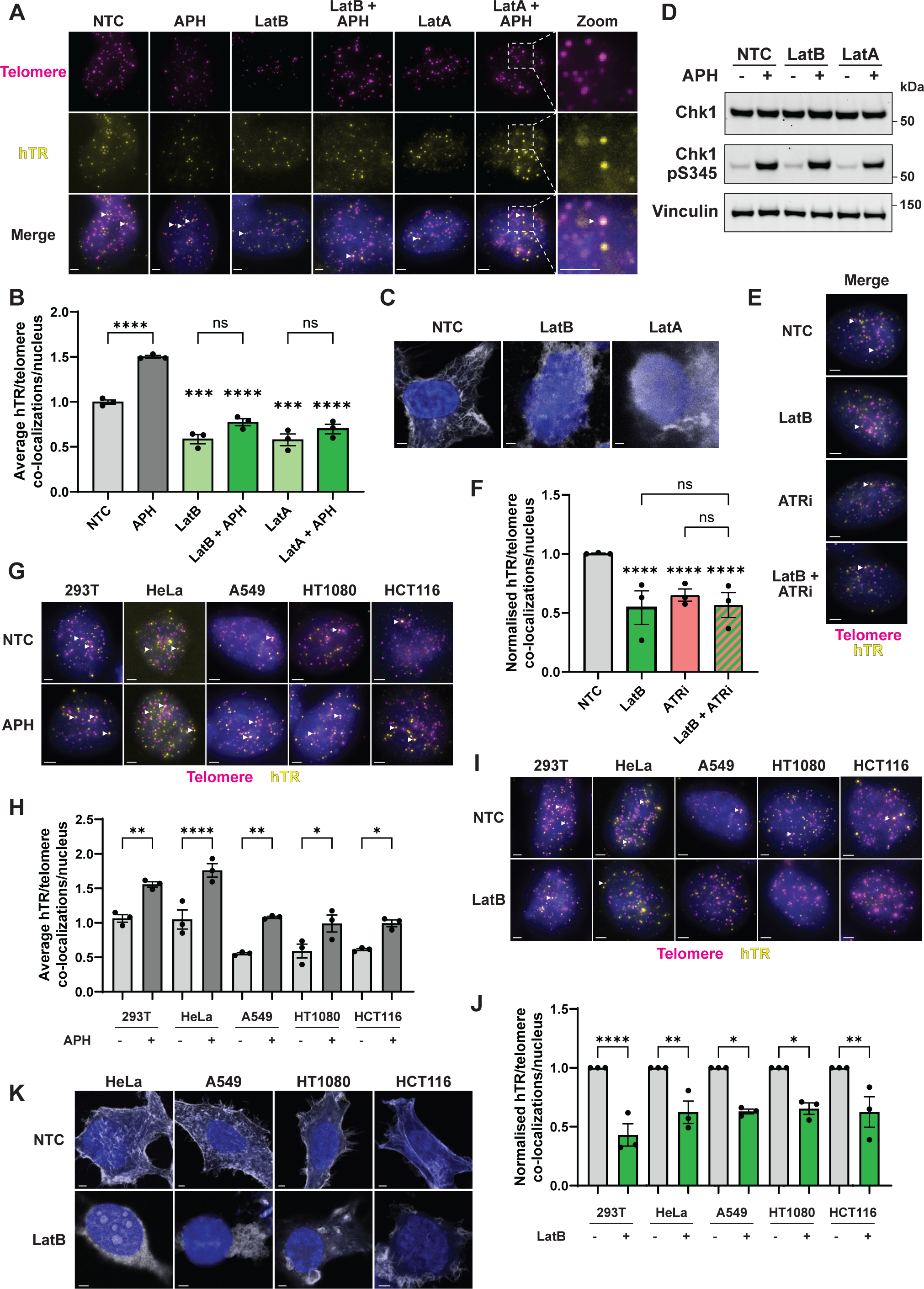
F-actin polymerization facilitates telomerase recruitment to telomeres. (A) Representative images of SABER FISH probing for telomeres (pink) and hTR (yellow) in 293T cells stained with DAPI (blue). Cells were treated with actin polymerization inhibitors LatB (0.2 μM) or LatA (50 nM), or DMSO (no treatment control, NTC), ± 1.5 μM APH. Co-localizations are indicated by white arrows in the merge row. Scale bar, 2 μm. (B) Average hTR/telomere co-localizations in S phase 293T cells following treatment with DMSO (NTC), LatB or LatA, ± APH. Significance for the latrunculin treated samples is expressed relative to the appropriate control (NTC ± APH). (C) Phalloidin staining (white) of 293T cells treated with DMSO (NTC), 0.2 μM LatB or 50 nM LatA also stained with DAPI (blue) and imaged by super-resolution Airyscan microscopy. Scale bar, 2 μm. (D) Western blot of 293T cells treated with DMSO (NTC), LatB or LatA, ± APH, probed for Chk1 and Chk1 pS345, with vinculin as a control. (E) SABER FISH for hTR (yellow) and telomeres (pink) with DAPI staining (blue) in 293T cells following treatment with DMSO (NTC), 0.2 μM LatB and 0.25 μM ATR inhibitor VE-822 (ATRi). Co-localizations are indicated by white arrows. Scale bar, 2 μm. (F) Normalized recruitment of telomerase to telomeres in 293T cells treated with DMSO (NTC), LatB, VE-822 (ATRi) or both LatB and ATRi. Significance for each column is expressed relative to the untreated control (NTC). (G) Representative images of SABER FISH for hTR (yellow) and telomeres (pink) with DAPI staining (blue) of nuclei in 293T, HeLa, A549, HT1080 and HCT116 cells treated with DMSO (NTC) or 1.5 μM APH. Co-localizations are indicated by white arrows. Scale bar, 2 μm. (H) Quantitation of telomerase recruitment in 293T, HeLa, A549, HT1080 and HCT116 cells ± APH. (I) SABER FISH for hTR (yellow) and telomeres (pink) with DAPI staining (blue) in 293T, HeLa, A549, HT1080 and HCT116 cells treated with 0.2 μM LatB. Co-localizations are indicated by white arrows. Scale bar, 2 μm. (J) Normalized quantitation of telomerase recruitment to telomeres from (I). Cells were treated with DMSO or 0.2 μM LatB. (K) Phalloidin (white) and DAPI (blue) staining of HeLa, A549, HT1080 and HCT116 cells treated with DMSO (NTC) or 0.2 μM LatB, imaged by super-resolution Airyscan microscopy. Scale bar, 2 μm. All bar graphs displayed as mean ± SEM; n = 3; ****p<0.0001, ***p<0.001,**p<0.01,*p<0.05. See also Figure S1.

This result suggests that F-actin is acting downstream of the DNA repair machinery that acts at stalled replication forks. To explore this, 293T cells were treated with an ATR kinase inhibitor (VE-822)^46^ and LatB, separately and in combination. The cells did not display considerably altered cell cycle progression due to treatment with these inhibitors (Figure S1D). While LatB treatment did not impact Chk1 phosphorylation, both treatments inhibited actin polymerization (Figures S1E-S1F). Both inhibitors induced a significant reduction in telomerase recruitment; however, simultaneous treatment with both did not result in a further reduction (Figures 1E-1F). These data suggest that F-actin functions downstream of ATR within the same pathway to mediate telomerase recruitment.

To validate the generality of this pathway, the requirement of ATR for recruitment was first confirmed in the cervical cancer cell line HeLa, as this has only been examined in 293T cells previously^8^. Knockdown of ATR by siRNA and ATR inhibition using VE-822 both resulted in a significant reduction in telomerase recruitment in HeLa cells, which was not accompanied by an observable perturbation to cell cycle profiles (Figures S1G-N). Subsequently, the levels of telomerase recruitment in response to replication stress was examined in a panel of telomerase positive human cell lines, including HeLa, lung adenocarcinoma A549, fibrosarcoma HT1080 and colorectal carcinoma HCT116 cells. Following APH treatment to induce replication stress, all cell lines displayed a significant increase in telomerase recruitment with no substantial alteration to their cell cycle profile (Figures 1G-1H, Figures S1O-P). Treatment of this panel of cell lines with LatB to inhibit actin polymerization also recapitulated the reduction in telomerase recruitment observed in 293T cells (Figures 1I-1K, Figure S1Q). Overall, these results provide evidence that the polymerization of actin in telomerase positive human cells is important for telomerase recruitment, and that replication stress-induced recruitment requires ATR-mediated F-actin formation.

### Telomerase recruitment is facilitated by known regulators of nuclear F-actin

Nuclear actin polymerization and function is controlled and regulated by a wide array of proteins. Monomeric actin is transported into the nucleus by importin 9; this transport is regulated by cofilin 1^47,48^. Both nuclear and cytoplasmic actin fibres can be nucleated by formin 2, and branching of the fibres is mediated by ARP2/3, which is regulated by WASP through mTOR^39^. F-actin depolymerization is necessary for fibre reorganization, and can be performed by cofilin 1, which is regulated by LIM kinase^49^. Furthermore, myosin motor proteins aid F-actin function by transporting cargo and aid remodelling of the F-actin network^50^.

To confirm that actin-mediated telomerase recruitment utilizes these known regulatory elements, 293T cells were treated with small molecule inhibitors of five key actin regulators: the mTOR inhibitor INK128^51^, the ARP2/3 inhibitor CK-666^52^, the WASP inhibitor wiskostatin^53^, the LIMK inhibitor LIMKi 3^54^, and the myosin inhibitors BDM (2,3-butanedione monoxime)^55^ and BTS (N-benzyl-p-toluene sulphonamide)^56^. Chemical inhibition of these actin regulators resulted in significantly decreased telomerase presence at telomeres (Figures 2A-2B, Figures S2A-C). The cells were also treated with APH briefly to induce replicative stress; while inhibition of these regulatory elements did not impair Chk1 phosphorylation (Figure 2C) or cell cycle progression (Figure S2A), all inhibitors prevented the increase in telomerase recruitment induced by replication stress (Figures 2A-2B, Figures S2B-C).

**Figure 2.**
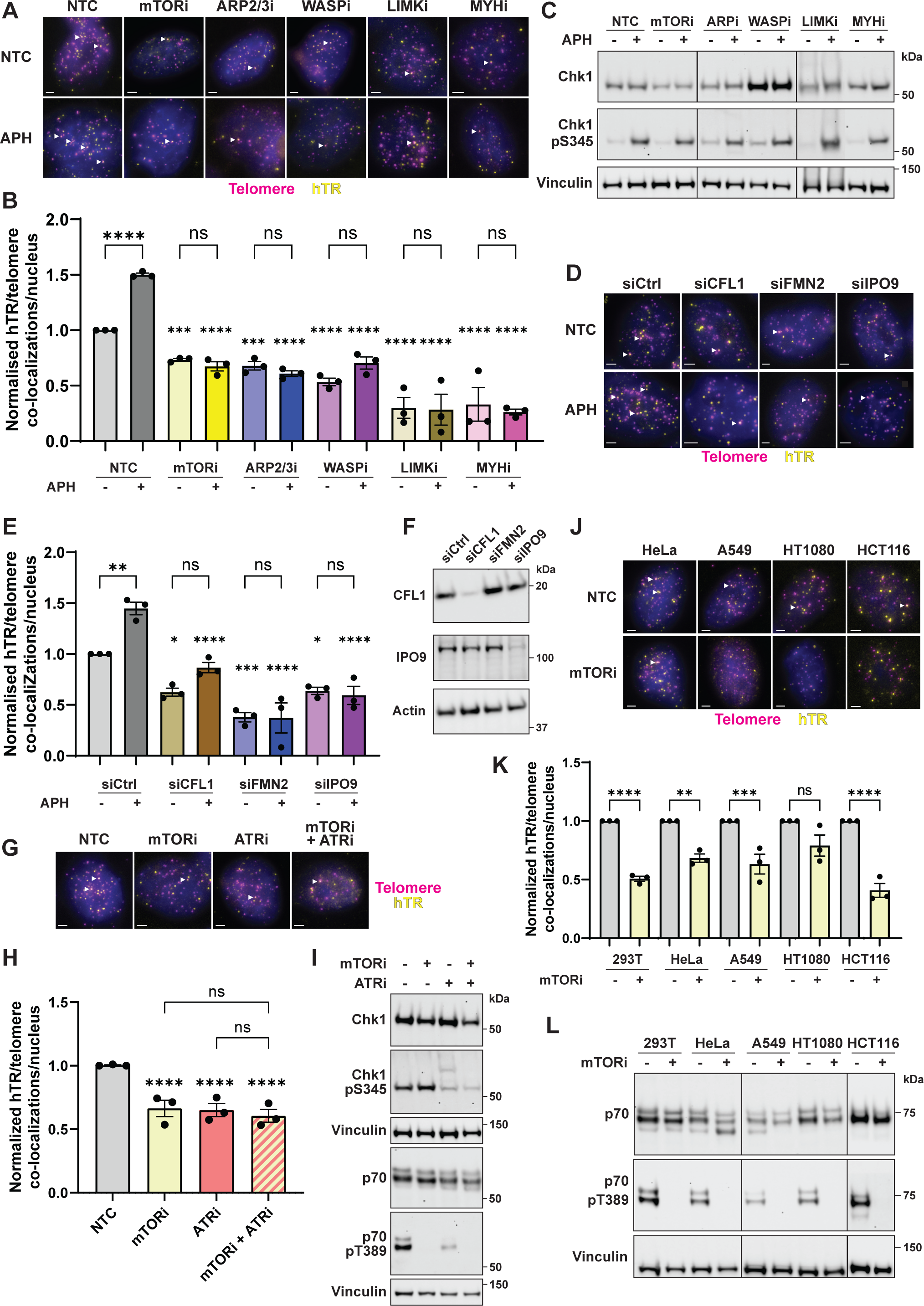
Telomerase recruitment is facilitated by known regulators of nuclear F-actin. (A) Representative SABER FISH images of 293T cells probed for hTR (yellow) and telomeres (pink) with DAPI (blue) staining. Cells were treated with DMSO (NTC), 0.2 μM INK128 (mTORi), 200 μM CK-666 (ARP2/3i), 5 μM wiskostatin (WASPi), 10 μM LIMKi-3 (LIMKi) or 10 mM BDM (MYHi), ± 1.5 μM APH. Co-localizations are indicated by white arrows. Scale bar, 2 μm. (B) Normalized hTR/telomere co-localizations in 293T cells treated with DMSO (NTC), INK128 (mTORi), CK-666 (ARP2/3i), wiskostatin (WASPi), LIMKi 3 (LIMKi) or BDM (MYHi), ± APH. Significance for each column is expressed relative to the appropriate control (NTC ± APH). (C) Western blots of 293T cells treated with DMSO (NTC), INK128 (mTORi), CK-666 (ARPi), wiskostatin (WASPi), LIMKi-3 (LIMKi) or BDM (MYHi), ± APH, probing for Chk1, Chk1 pS345 and vinculin as a control. (D) Representative SABER FISH images of 293T cells transfected with control, cofilin 1 (CFL1), formin 2 (FMN2) or importin 9 (IPO9) siRNA, ± APH treatment. Cells were stained with DAPI (blue) and probed for hTR (yellow) and telomeres (pink). Co-localizations are indicated by white arrows. Scale bar, 2 μm. (E) Normalized telomerase presence at telomeres in 293T cells transfected with control, cofilin 1 (CFL1), formin 2 (FMN2) or importin 9 (IPO9) siRNA, ± 1.5 μM APH treatment. Significance for each siRNA treated sample is expressed relative to the appropriate control (siCtrl ± APH). (F) Western blot of 293Tcells transfected with control, CFL1, FMN2 or IPO9 siRNA, probing for CFL1, IPO9 and actin as a control. Note: FMN2 western blot not shown due to lack of specific antibody. (G) Representative FISH images of 293T cells treated with DMSO (NTC), 0.2 μM INK128 (mTORi) or 0.25 μM VE-822 (ATRi) alone and in combination. Cells were stained with DAPI (blue) and probed for hTR (yellow) and telomeres (pink). Co-localizations are indicated by white arrows. Scale bar, 2 μm. (H) Normalized hTR/telomere co-localizations in 293T cells treated with DMSO (NTC), INK128 (mTORi), VE-822 (ATRi), or mTORi and ATRi together. Significance for each column is expressed relative to the untreated control (NTC). NTC and ATRi data are the same as presented in Figure 1F. (I) Western blotting of 293T cells treated with DMSO, INK128 (mTORi), VE-822 (ATRi), or mTORi and ATRi in combination. Blots were probed for Chk1, Chk1 pS345, p70, p70 pT389 (an mTOR substrate) and vinculin as a loading control. (J) SABER FISH images of HeLa, A549, HT1080 and HCT116 cells treated with DMSO (NTC) or 0.2 μM INK128 (mTORi), probing for hTR (yellow) and telomeres (pink) with DAPI staining (blue). Co-localizations are indicated by white arrows. Scale bar, 2 μm. (K) Normalized telomerase presence at telomeres in 293T, HeLa, A549, HT1080 and HCT116 cells treated with DMSO or INK128 (mTORi). (L) Western blot of 293T, HeLa, A549, HT1080 and HCT116 cells treated with DMSO or INK128 (mTORi), probing for p70, p70 pT389 and vinculin as a loading control. All bar graphs displayed as mean ± SEM; n = 3; ****p<0.0001, ***p<0.001,**p<0.01,*p<0.05. See also Figure S2.

We next tested the role of several regulators of nuclear actin polymerization by siRNA knockdown in 293T cells; this included cofilin 1 (encoded by the gene *CFL1*), formin 2 (*FMN2*) and importin 9 (*IPO9*). Knockdown of these proteins resulted in a significant decrease in telomerase recruitment (Figures 2D-2F, Figures S2D-F). Furthermore, cells briefly treated with APH following knockdown retained a functional DNA damage response (Figure S2G) and normal cell cycle profile (Figure S2H), but again showed no increase in telomerase recruitment upon replication stress (Figures 2D-E, Figures S2D-E). Overall, these data demonstrate that the increased telomerase recruitment induced by replication stress requires a functional regulatory system to promote nuclear import and polymerization of actin.

Given that mTOR is phosphorylated in an ATR-dependent manner in response to replication stress and is necessary for replication stress-induced nuclear actin polymerization^34^, we next validated that this axis is required for actin-mediated telomerase recruitment. 293T cells were treated with VE-822 and INK128, separately and in combination, to assess the impact on recruitment. Neither treatment drastically perturbed cell cycle progression (Figure S2I). While both inhibitors significantly decreased telomerase recruitment as expected, dual treatment did not further reduce telomerase presence at telomeres (Figures 2G-I). These data indicate that the ATR-mTOR kinase pathway regulates telomerase recruitment via F-actin polymerization.

To establish the universality of this pathway, the selected panel of telomerase positive cell lines (HeLa, A549, HT1080 and HCT116) was treated with INK128 to assess its effect on telomerase recruitment. Following mTOR inhibition, all cell lines displayed decreased telomerase recruitment without major changes to cell cycle progression (Figures 2J-L, Figure S2J). Overall, the results from these experiments indicate that the known nuclear actin regulatory network is important for facilitating telomerase localization to telomeres, both under endogenous conditions and following induction of exogenous replication stress.

### Nuclear F-actin polymerization is required for telomerase recruitment

The role of nuclear F-actin in resolving replication stress^34,37^, in combination with our observation that knockdown of IPO9 impacted telomerase recruitment (Figure 2E, Figure S2E), suggested that it is the nuclear pool of F-actin that specifically regulates telomerase recruitment, rather than an indirect effect of perturbations to cytoplasmic actin. To directly test this hypothesis, we utilized expression constructs encoding wild type (WT) actin or a polymerization-deficient actin mutant (R62D) tagged with a nuclear localization signal (NLS)^32,34^. Both constructs, along with an empty vector as a control, were transfected into 293T cells, where only R62D actin disrupted the polymerization of nuclear F-actin (Figure 3A, Figures S3A-B). Expression of exogenous WT actin did not impact telomerase recruitment under endogenous or stressed conditions; however, expression of the R62D mutant significantly decreased recruitment in both conditions (Figures 3B-D). To further validate this result, expression of WT and R62D actin was then repeated with simultaneous siRNA knockdown of IPO9 to reduce the nuclear pool of endogenous actin (Figure 3E, Figures S3C-D). IPO9 knockdown reduced telomerase recruitment in cells transfected with empty vector as expected; however, expression of exogenous WT actin was able to rescue this phenotype by replenishing the nuclear actin pool (Figures 3F-G). Conversely, expression of the mutant R62D mutant significantly reduced telomerase recruitment, regardless of IPO9 knockdown. These data provide evidence that the regulation of telomerase recruitment specifically involves nuclear F-actin filaments.

**Figure 3.**
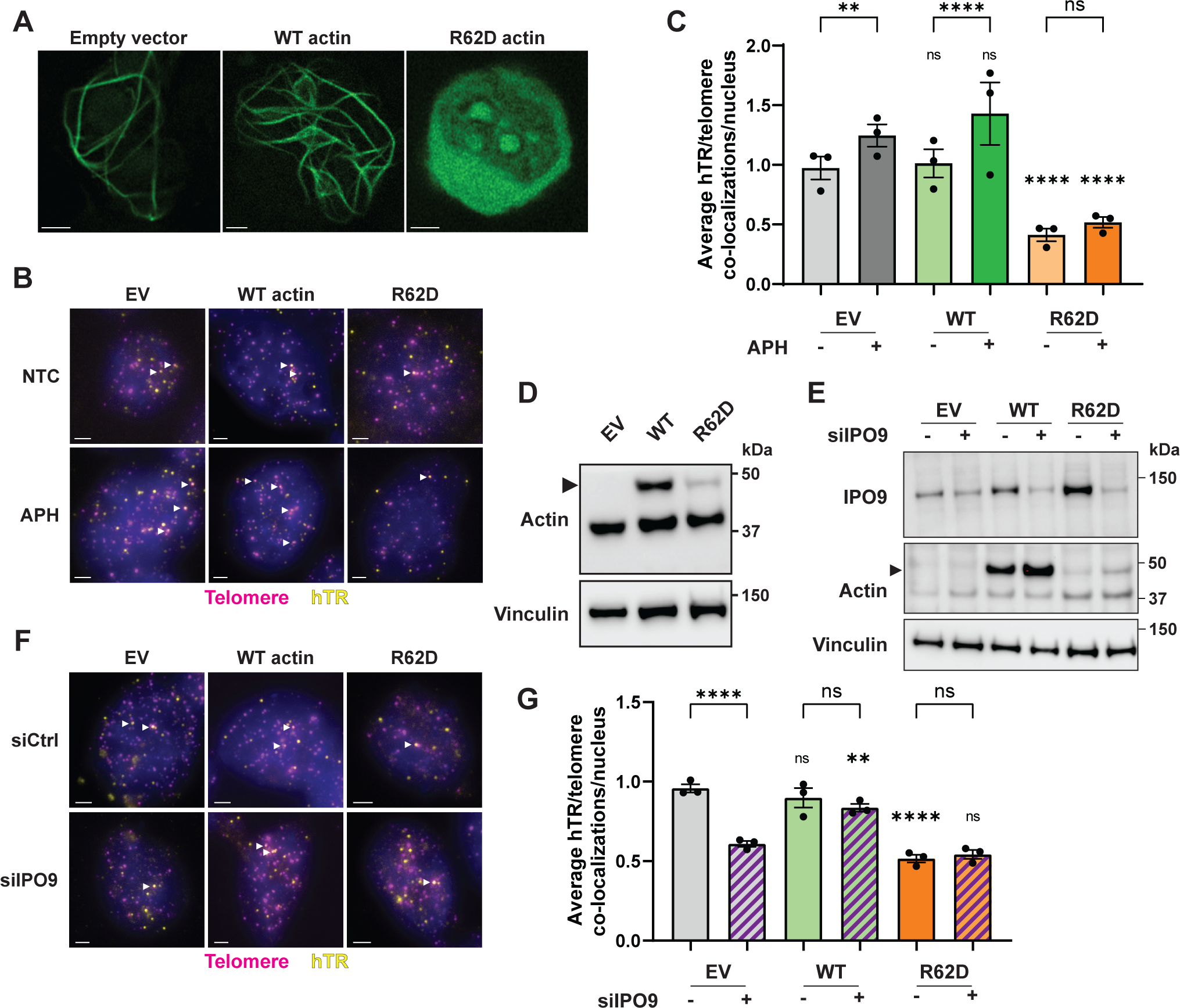
Nuclear F-actin polymerization is required for telomerase recruitment. (A) Super-resolution Airyscan microscopy of 293T cells transfected with vectors encoding WT or polymerization-deficient mutant (R62D) actin, using empty vector as a control. Cells were co-transfected with a BFP- and NLS-tagged actin chromobody (nuclear-actin-ChB) to visualize nuclear F-actin (green). Scale bar, 2 μm. (B) Representative images of 293T cells probed for hTR (yellow) and telomeres (pink) with DAPI (blue) staining following transfection with vectors encoding WT or mutant (R62D) actin and treated with DMSO or 1.5 μM APH. Empty vector (EV) was used as a control. Co-localizations are indicated by white arrows. Scale bar, 2 μm. (C) Average telomerase presence at telomeres in 293T cells transfected with empty vector (EV) or vector encoding WT or mutant (R62D) actin, treated with DMSO or APH. Significance for each sample is expressed relative to the appropriate control (EV ± APH). (D) Western blot of 293T cells transfected with empty vector (EV) or vector encoding WT or mutant (R62D) actin, probing for actin and vinculin as a control. Exogenous actin is tagged with a 3×NLS, and is indicated by a black arrow. (E) Western blot of 293T cells transfected with vectors encoding WT or mutant (R62D) actin, with or without IPO9 siRNA. Empty vector (EV) and control siRNA were used as controls. Blots were probed for IPO9, actin and vinculin as a loading control. Exogenous actin is tagged with a 3×NLS, and is indicated by a black arrow. (F) Representative FISH images of 293T cells probed for hTR (yellow) and telomeres (pink) with DAPI (blue) staining. Cells were transfected with empty vector (EV) or vectors encoding WT or mutant (R62D) actin together with control or IPO9 siRNA. Co-localizations are indicated by white arrows. Scale bar, 2 μm. (G) Average hTR/telomere co-localizations in 293T cells transfected with vectors encoding WT or mutant (R62D) actin, with or without IPO9 siRNA. Empty vector (EV) and control siRNA were used as controls. Significance for each sample is expressed relative to the appropriate control (EV ± siIPO9). All bar graphs displayed as mean ± SEM; n = 3; ****p<0.0001, **p<0.01. See also Figure S3.

### Telomerase recruitment occurs closer to the nuclear periphery under replication stress conditions

One of the emerging functions of nuclear F-actin in resolving genomic stress is the re-localization of stalled replication forks or sites of DNA damage to the nuclear periphery^33,34,36^. It has also been demonstrated that telomeres undergoing replicative stress are repositioned toward the nuclear periphery in an F-actin dependent manner^34,38^. Given this, we hypothesized that telomere re-localization may be required prior to telomerase recruitment. To investigate this, the nuclear periphery was first identified by DAPI staining of 293T cells. Individual hTR or telomere foci, as visualized by FISH, were then identified alongside co-localized hTR/telomere foci indicating telomerase recruitment (Figure 4A). The minimum distance between each focus and the nuclear periphery was then calculated. There was no significant difference in localization between hTR or telomeres; however, telomerase interacting with telomeres was slightly, but significantly, further from the nuclear periphery than telomeres or hTR alone (Figure 4B, Figure S4A). This suggests that while the nuclear periphery serves as a site for DNA repair, it is not where telomerase recruitment predominately occurs under basal conditions.

**Figure 4.**
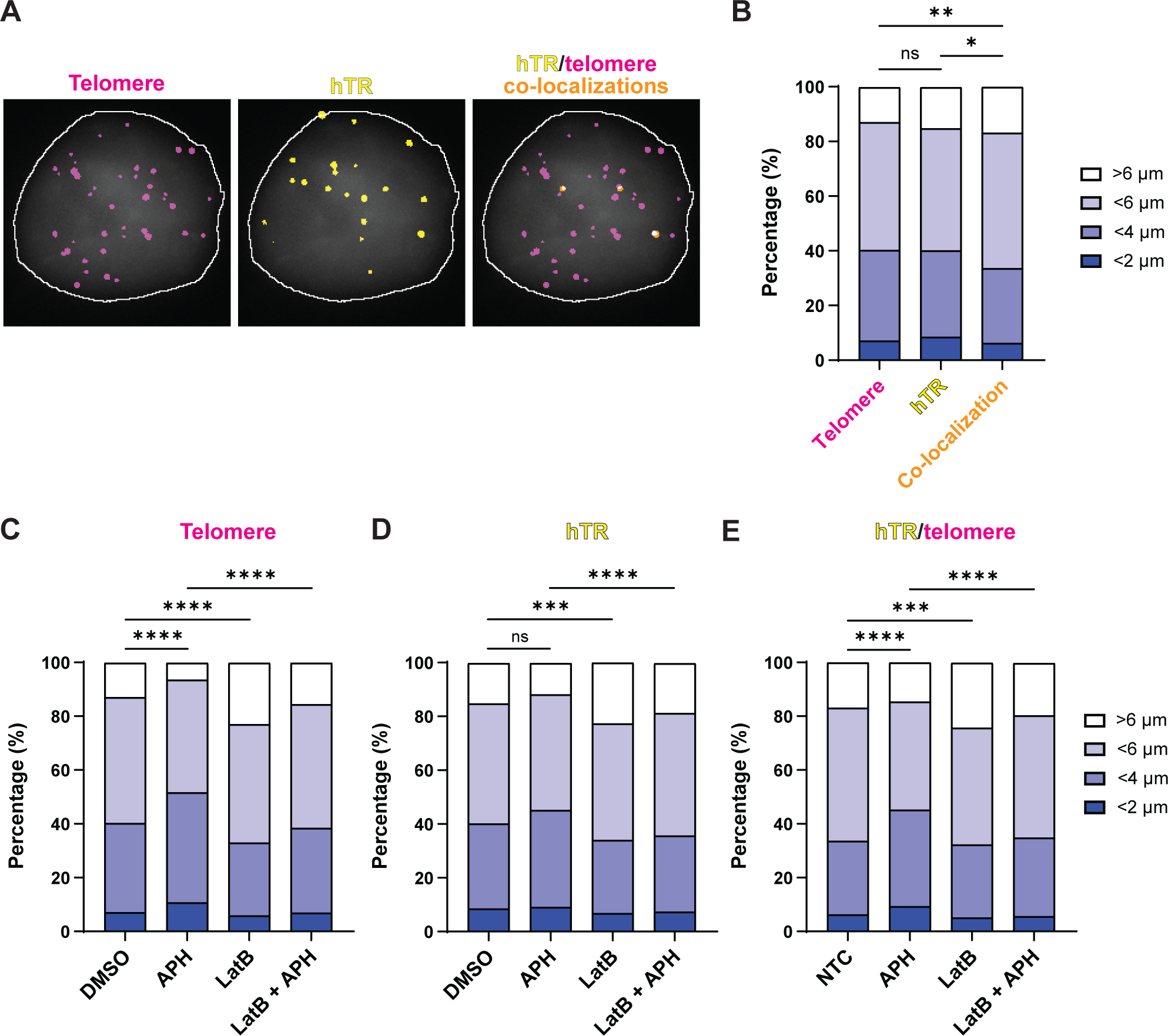
Telomeres interacting with telomerase move closer to the nuclear periphery under replication stress conditions. (A) Representative images showing identified telomere (pink), hTR (yellow) and co-localizing hTR/telomere (orange) foci. 293T nuclei were stained using DAPI (grey) to identify the nuclear periphery (white outline). Nuclei, foci and co-localizations were detected using CellProfiler. (B) Minimum distance of telomeres, hTR, or hTR/telomere co-localizations from the nuclear periphery in 293T cells. Data are displayed as percentages based on distance from nuclear periphery. Statistics were performed using *χ*^2^ tests. n = 14756 (telomeres), 6123 (hTR) and 624 (co-localizations). (C-E) Minimum distance of telomeres (C), hTR (D), or hTR/telomere co-localizations (E) from the nuclear periphery in 293T cells treated ± 1.5 μM APH and 0.2 μM LatB. Data are displayed as percentages based on distance from nuclear periphery. Statistics were performed using *χ*^2^ tests. NTC data are the same as presented in Figure 4B. Telomeres: n = 14756 (NTC), 19360 (APH), 11030 (LatB) and 12708 (LatB + APH). hTR: n = 6123 (NTC), 8981 (APH), 9839 (LatB) and 7207 (LatB + APH). Co-localizations: n = 624 (NTC), 910 (APH), 988 (LatB) and 864 (LatB + APH). ****p<0.001, ***p<0.001,**p<0.01,*p<0.05. See also Figure S4.

The positioning of telomerase and telomeres in the nucleus was also measured following APH treatment. Individual telomeres were on average closer to the nuclear periphery following induction of replication stress (Figure 4C, Figure S4B). This re-localization was F-actin mediated, as telomeres in LatB treated cells were further from the periphery than in untreated cells and did not reposition towards the periphery following APH treatment (Figure 4C, Figure S4B). This trend is also seen with individual hTR foci and those which co-localize with telomeres (Figures 4D-E, Figures S4C-D). Overall, these data suggest that telomeres, both with and without co-localized telomerase, are sensitive to replication stress and are consequently moved closer to the nuclear periphery. Furthermore, nuclear F-actin facilitates reorganization of the nucleus, as telomeres and telomere-telomerase co-localizations are further from the periphery in cells with F-actin inhibition.

### Telomerase recruitment to telomeres occurs in proximity to F-actin filaments

We hypothesized that F-actin might also serve as a direct site of interaction between telomeres and telomerase. To test this, endogenous nuclear actin was visualized using an actin chromobody^57^, tagged with both blue fluorescent protein (BFP) and an NLS (nuclear-actin-ChB). FISH against hTR and telomeres was then performed on HeLa cells expressing nuclear-actin-ChB and treated briefly with APH to promote F-actin formation. Neither nuclear-actin-ChB expression nor APH treatment impacted the cell cycle progression of the BFP-positive cells (Figure S5A). Cells were visualized by confocal laser scanning microscopy using an Airyscan super-resolution detector (Carl Zeiss), where co-localized hTR/telomere foci were observed residing on F-actin (Figures 5A-5B). To quantify this, individual and co-localizing hTR/telomere foci were identified along with F-actin, and the minimum distance between each focus and the nearest actin fibre was then calculated. While the mean distance of hTR and telomere foci from actin fibres were not significantly different, telomerase foci co-localizing with telomeres were on average significantly closer to F-actin than either unbound telomerase or telomeres (Figure 5C). This result was due to more foci residing directly on or adjacent to F-actin, as well as fewer foci which were distant (> 1 μm) from F-actin (Figures S5B-S5C).

**Figure 5.**
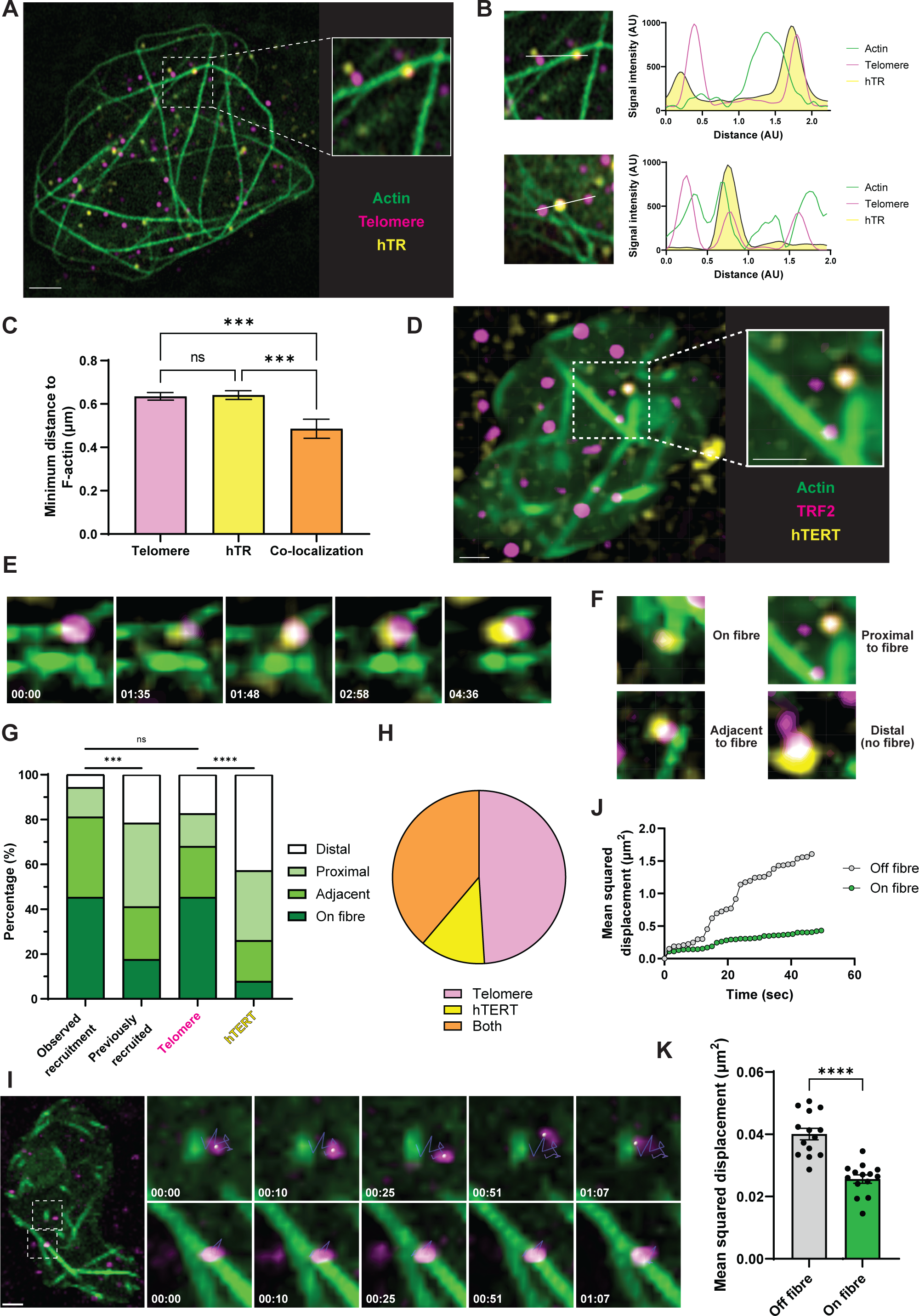
Telomerase recruitment to telomeres occurs in proximity to F-actin. (A) Representative image of HeLa cells expressing nuclear-actin-ChB to visualize nuclear F-actin (green), with telomere (pink) and hTR (yellow) staining by SABER FISH, imaged via super-resolution Airyscan microscopy. Scale bar, 2 μm. (B) Line scan of telomere (pink) and hTR (yellow) signals to demonstrate co-localization along nuclear F-actin (green). (C) Mean minimum distance of telomeres, hTR and hTR/telomere co-localizations from nuclear F-actin. Data are pooled from >100 nuclei; n = 7384 (telomeres), 5929 (hTR) and 434 (co-localizations). (D) Still image from live cell imaging experiments. Nuclear-actin-ChB to visualize nuclear F-actin (green) was transfected into CRISPR-modified HeLa cells expressing HA-mEOS3.2-tagged TRF2 (pink) and FLAG-HaloTag-tagged hTERT (visualized using JF-646-HaloTag ligand; yellow). Scale bar, 2 μm. (E) Time course of live cell imaging experiment showing hTERT (yellow) and TRF2 (pink) foci co-localizing on a nuclear F-actin fibre (green), over the indicated time in minutes. (F) Classification used for live cell imaging quantification. (G) Quantification of live cell imaging experiments, displaying distribution of foci with respect to F-actin. ‘Observed recruitment’ includes co-localizations where one or both hTERT/TRF2 foci were identified alone prior to recruitment. ‘Previously recruited’ includes co-localizations where both foci were identified simultaneously. Quantified telomere (TRF2) and hTERT foci did not co-localize with the other. Statistics were performed using *χ*^2^ tests. n = 52 (observed recruitment), 51 (previously recruited), 247 (telomeres) and 36 (hTERT). (H) Observed recruitment events (Figure 5G; excluding distal events) represented based on closest constituent focus to nuclear F-actin (n = 49). (I) Time course of live cell imaging experiment over the indicated time in minutes, showing telomere (TRF2; pink) foci mobility in relation to F-actin (green). Top panels display a mobile telomere which is off F-actin, while the bottom panels show a static telomere on F-actin. Scale bar, 2 μm. (J) Example cumulative mean squared displacement (MSD) of telomere (TRF2) foci in relation to F-actin. MSD is plotted over time for a focus which is not on F-actin (grey) or one that remains associated with F-actin (green). (K) MSD of telomere (TRF2) foci from live cell imaging experiments. MSD was calculated for each telomere on a frame-by-frame basis, and each MSD value was classified as being “Off fibre” or “On fibre” based on the intensity of nuclear-actin-ChB signal within the focus region. Data are displayed as means from 14 experiments, representing >63 nuclei and >25,000 MSD values. All bar graphs displayed as mean ± SEM; ****p<0.0001, ***p<0.001. See also Figure S5, Videos S1-3.

To explore this further, live cell imaging was performed using CRISPR-modified HeLa cells expressing endogenous hTERT tagged with a FLAG-HaloTag and endogenous TRF2 tagged with the fluorescent protein HA-mEOS3.2, to allow visualization of telomerase and telomeres, respectively^58^. These cells were transfected with the nuclear-actin-ChB, synchronized by double thymidine block (Figure S5D), and released into S phase for 3-5 hrs prior to live cell imaging using super-resolution microscopy (Figure 5D, Videos S1 and S2). Synchronization by double thymidine block also promotes formation of nuclear F-actin. Telomerase is known to rapidly bind and dissociate from telomeres in S phase, while longer interactions with telomeres are rarer^31,58^. These longer interactions presumably signify successful recruitment of telomerase leading to productive telomere extension. Given this, we assessed these long telomerase/telomere interactions to determine whether proximity to F-actin would help mediate this process. Stable interactions (> 30 sec) between telomerase and telomeres (Figure 5E) were classified based on where they occurred relative to F-actin; this included on or adjacent to a fibre, or in proximity to F-actin (∼1 μm maximum distance), with all other events classified as distal (Figure 5F). Stable interactions were also separately classified based on whether recruitment had already occurred prior to imaging, or if the recruitment was observed occurring (i.e. at least one focus was observed prior to it co-localizing with its partner). Using these classifications, ∼45% of observed recruitment events occurred directly on F-actin, with ∼35% of the remaining events occurring adjacently (Figure 5G). By comparison, previous recruitment events were imaged directly on or adjacent to F-actin only ∼15% and ∼25% of the time, respectively. The distribution of these events was significantly different, which suggests that recruitment may occur in proximity to F-actin, but this is not required for the continued interaction between telomerase and a telomere.

It was also observed by live cell imaging that telomeres displayed a similar profile to observed recruitment events; a majority of telomeres which were never bound by telomerase were on or adjacent to F-actin (Figure 5G). In contrast, telomerase foci which never interacted with a telomere were primarily located distally from F-actin. This suggested that under replication stress conditions, telomeres are the first to interact with actin filaments, which ultimately could facilitate the ability of telomerase to interact with those telomeres. When the position of the telomere and telomerase foci relative to actin fibres was examined, in ∼50% of observed recruitment events the telomere was located closer to F-actin than telomerase, while telomerase was closer only ∼10% of the time (the remaining ∼40% were equidistant) (Figure 5H).

We observed that telomeres located on F-actin were less mobile than those distant from fibres (Figure 5I, Video S3); this was quantified by measuring the mean squared displacement (MSD) of each telomere on a frame-by-frame basis and classifying each focus as “on” or “off” F-actin based on the intensity of nuclear-actin-ChB signal within the focus (examples in Figure 5J, Figures S5E-F). From this analysis, telomeres which were on F-actin were ∼40% less mobile than other telomeres (Figure 5K, Figure S5G).

Overall, these data suggest that under replicative stress, telomeres are restrained by F-actin and that these telomeres may serve as a site for telomerase recruitment to facilitate telomere elongation.

### Stalled replication forks promote telomerase recruitment at F-actin

While it is known that replication stress promotes telomerase recruitment^8^, the exact mechanism behind this process remains unclear. One possible explanation for this phenomenon is that replication fork stalling within a telomere triggers telomerase recruitment to that same telomere. To examine the hypothesis that telomerase recruitment occurs in proximity to stalled replication forks, PCNA (proliferating cell nuclear antigen) foci were used as a marker of DNA replication^59^. Immunofluorescence (IF) against PCNA was combined with SABER FISH for hTR and telomeres (Figure 6A) in 293T cells in the presence or absence of APH. While a basal level of co-localization between all three elements was observed in untreated cells, APH treatment resulted in ∼4-fold more hTR-telomere co-localizations that overlapped with PCNA foci (Figure 6B). As expected, APH treatment also increased telomere-PCNA or hTR-telomere co-localizations ∼2-fold (Figures 6C-6D). Interestingly, when hTR-telomere co-localizations were analyzed according to their overlap with PCNA, the proportion of recruitment events which coincided with PCNA approximately doubled following APH treatment (Figure 6E, Figure S6A).

**Figure 6.**
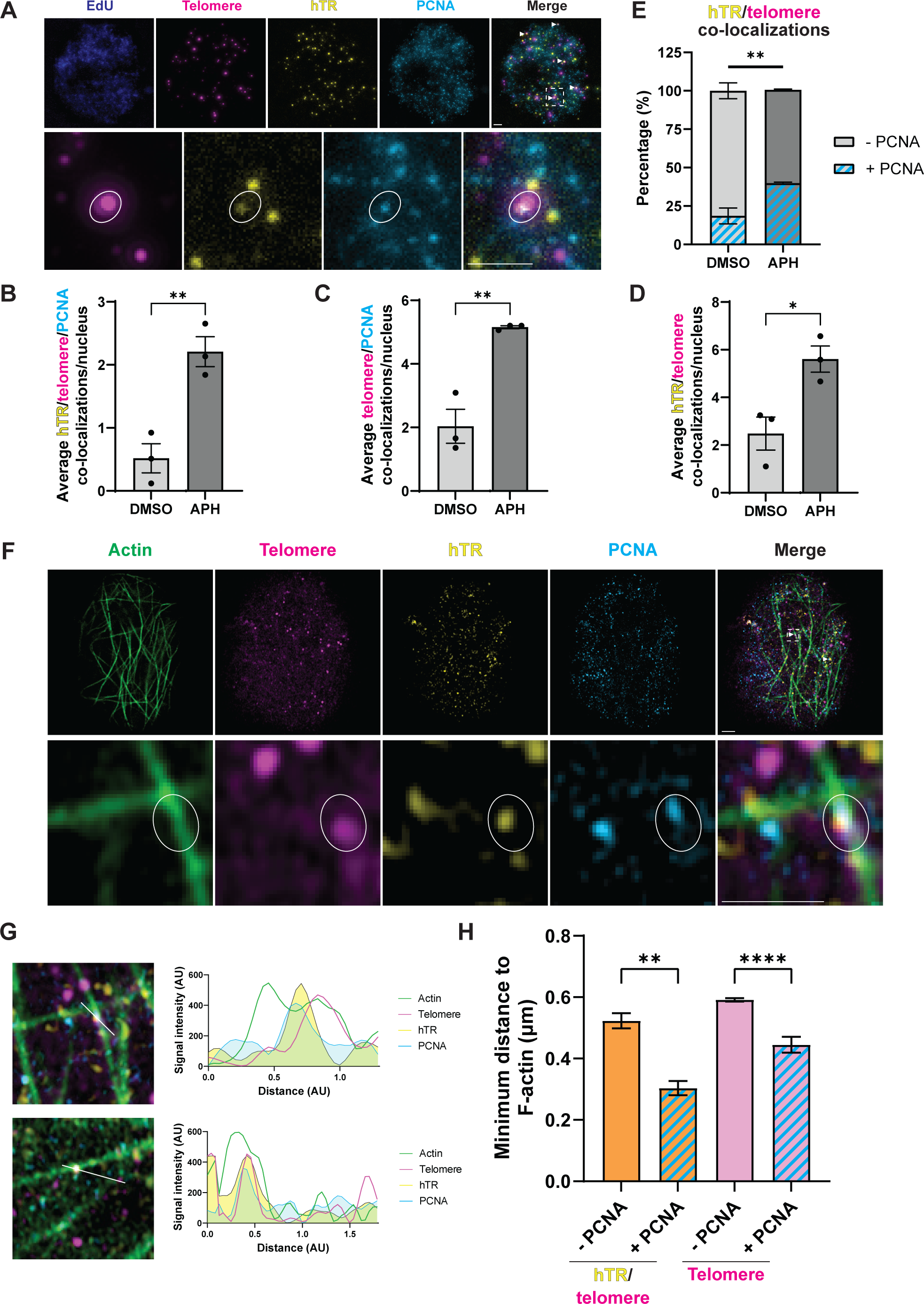
Stalled replication forks promote telomerase recruitment at F-actin. (A) Representative image of 293T cells stained using SABER FISH probes against telomeres (pink) and hTR (yellow) with immunofluorescence against PCNA (cyan). S phase cells were identified with EdU (blue). Co-localizations between hTR/telomere/PCNA are indicated by white arrows in the merge panel and circles in the zoomed panels. Scale bars, 2 μm. (B-D) Average co-localizations between hTR/telomere/PCNA (B), telomere/PCNA (C) or hTR/telomere (D) in 293T cells ± APH. (E) Distribution of hTR/telomere co-localizations (shown in Figure 6D) which also co-localized with PCNA foci in 293T cells ± APH. (F) Representative image of HeLa cells transfected with nuclear-actin-ChB to visualize nuclear F-actin (green). Cells were stained via SABER IF-FISH to visualize telomeres (pink), hTR (yellow) and PCNA (cyan) before imaging using super-resolution Airyscan microscopy. Co-localization between hTR/telomere/PCNA is indicated by white arrows in the merge panel and a circle in the zoomed panels. Scale bars, 2 μm. (G) Line scan of telomere (pink), hTR (yellow) and PCNA (cyan) signals to demonstrate co-localization along nuclear F-actin (green). (H) Mean minimum distance of hTR/telomere co-localizations (left) and telomeres (right) from nuclear F-actin. Foci which overlapped with PCNA were compared to those which did not overlap. Data are pooled from 5 experiments, representing >200 nuclei; n = 562 (hTR/telomere), 171 (hTR/telomere/PCNA), 15098 (telomeres) and 418 (telomere/PCNA). All bar graphs displayed as mean ± SEM; n = 3 (all bar graphs except Figure 6H); ****p<0.0001, **p<0.01,*p<0.05. See also Figure S6.

Given the convergence between replication, telomerase recruitment and actin polymerization, we also performed SABER IF-FISH against PCNA, hTR and telomeres in HeLa cells expressing the nuclear-actin-ChB to assess the spatial relationship between these factors. Cells were briefly treated with APH to induce F-actin polymerization and imaged by super-resolution microscopy, in which hTR/telomere/PCNA co-localizations were observed residing on F-actin (Figures 6F-6G). Using the same strategy as described above, the minimum distance from F-actin for each focus alone and co-localizing with other foci was quantified. Both telomeres and hTR/telomere co-localizations which overlapped with PCNA were significantly closer to F-actin than those that did not overlap with PCNA (Figure 6H, Figures S6B-S6C), while there was no significant difference in the distance from F-actin between PCNA, telomerase or telomeres which did not co-localize with each other (Figures S6D-S6E). PCNA foci which reside along nuclear F-actin have been demonstrated to extensively co-localize with FANCD2^34^, indicating they are undergoing replication stress and may represent stalled replication forks^60^. Together these data indicate that F-actin directly facilitates telomerase recruitment to telomeres undergoing replication stress.

## DISCUSSION

In this study we have demonstrated that nuclear F-actin is necessary for the recruitment of human telomerase to telomeres. This process is regulated by the kinases ATR and mTOR, and relies upon a network of known actin regulators to facilitate nuclear actin polymerization. Telomeres are re-localized toward the nuclear periphery following replication stress, but their random movement around the nucleus is reduced by interaction with actin fibres. Telomerase interactions with telomeres predominately occur near stalled telomeric replication forks which are tethered to nuclear actin fibres. Overall, this suggests a tightly regulated process which governs the occurrence and timing of DNA replication, the replication stress response, and telomere maintenance. Our data support a model wherein telomeric replication stress, a natural occurrence in the highly repetitive telomere sequence^18,61,62^, triggers the DDR and nuclear F-actin polymerization (Figure 7A), ultimately resulting in telomere length maintenance. We hypothesize that this maintenance occurs via two mechanisms (Figures 7B-C) which are engaged depending on the extent of replication stress and whether telomeric replication forks can be restarted or are critically stressed.

**Figure 7.**
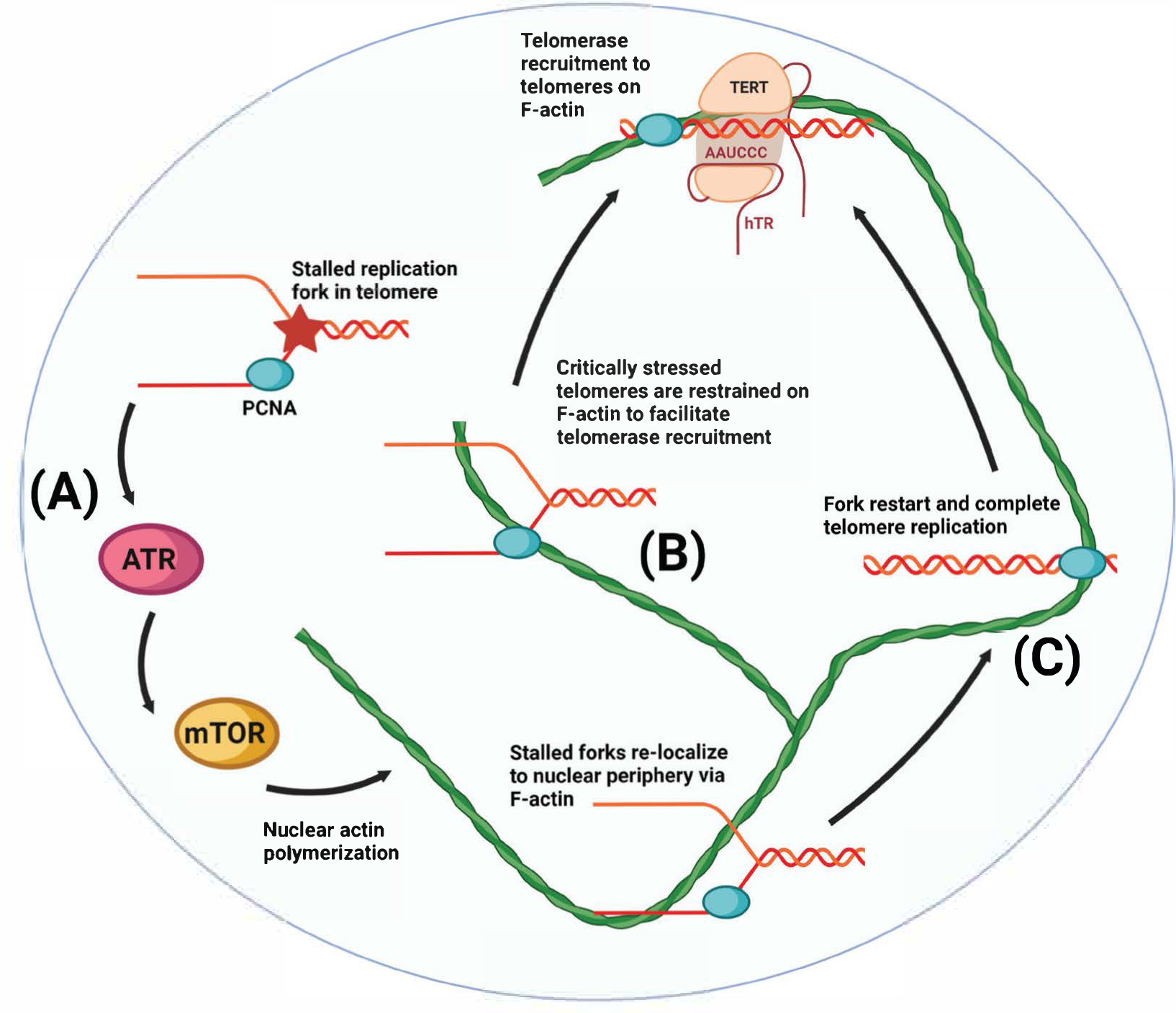
Models for replication- and actin-mediated telomerase recruitment. During S phase, DNA replication proceeds through telomeres which can result in fork stalling, activating ATR to facilitate restart or repair (A). ATR in turn activates mTOR to facilitate F-actin polymerization. (B) Telomeres which are critically stressed cannot be restarted and are restrained on F-actin to facilitate telomerase recruitment, resulting in the extension of the telomere to maintain its integrity. (C) Stalled telomeric replication forks which can be repaired are re-localized to the nuclear periphery via F-actin to enable fork restart. Once telomere replication has been completed, telomeres move away from the nuclear periphery where telomerase can be recruited in an F-actin-dependent manner. Created with Biorender.com.

Our data suggest that telomeres which are critically stressed through replication fork collapse are restrained by nuclear F-actin filaments to facilitate telomerase recruitment in lieu of successful telomere replication (Figure 7B). One potential outcome of replication stress is the sudden loss of telomeres^63–65^; a mechanism cancer cells may use to overcome this is targeting telomerase to those telomeres to ensure that telomere length is maintained. This is supported by the finding that, in yeast and human cells, telomerase expression aids cell survival following replication stress^66–68^. Telomerase can interact with telomeres through short ‘probing’ or long static interactions, and while we did not examine ‘probing’ telomerase molecules, other work has shown these molecules possess diffusive searching movement^31,58^. This could suggest that telomerase does not itself track along actin filaments to locate a critically stressed or shortened telomere, but instead that telomeres are restrained to facilitate encounters with telomerase. This is supported by our findings that telomeres residing on F-actin are less mobile than other telomeres (Figure 5I-K), and that telomerase recruitment predominately occurs in proximity to F-actin (Figure 5G).

Stalled replication forks move along actin filaments, which are required for the recovery of DNA replication after stalling^34^. We therefore hypothesize that telomeres which are mildly stressed are instead re-localized to the nuclear periphery to allow replication fork restart, complete telomere replication and subsequent telomerase recruitment (Figure 7C). This model is supported by other studies, as long-term treatment of cells with APH to induce mild replication stress ultimately results in telomere lengthening^18^. Expression of mutant POT1 (POT1-ΔOB), which cannot bind to the single-stranded telomeric overhang, results in telomere-specific replication stress, which causes telomere re-localization to the nuclear periphery^38^, telomerase recruitment^31^ and telomere extension^8,69^. Furthermore, depletion of mouse POT1b (which regulates single-stranded telomere overhang length) results in initial telomere shortening accompanied by a telomeric DDR which results in ATR-dependent telomerase recruitment^42,70^. However, long term growth of POT1b-deficient tumors ultimately causes telomere hyper-elongation in a telomerase-dependent manner^42^.

While chemically-induced replication stress using APH heightens recruitment, the inhibition of ATR^8^, actin polymerization or actin regulators (Figures 1A-B, 2A-B) result in a significant reduction in telomerase presence at telomeres under both endogenous and stressed conditions, which supports the notion that these are not two separate pathways, but one pathway which is overstimulated by exogenous induction of replication stress. While nuclear actin fibres are only occasionally detectable in unperturbed human cancer cell lines using the nuclear-actin-ChB (this study and ^34,37^), it has been demonstrated that ∼20% of unperturbed S phase cancer cells do contain nuclear actin filaments, detectable by immunofluorescence against tagged NLS-actin^37^. Elevated replication stress is a hallmark of cancer cells^71^, so it can be concluded that cancer cells likely utilize the same mechanisms to bring telomerase to telomeres under endogenous and exogenous stress-induced conditions.

From this study and previous work^38^, it appears that replication fork stalling within telomeres triggers telomere repositioning to the periphery via F-actin, which likely results in replication fork restart or activation of dormant origins of replication. Although it is unknown exactly how stalled telomeric replication forks might restart, it potentially involves the CST complex, which is well established to maintain telomeres^72^ but also aids in global replication fork restart^73^ through its ability to protect stalled forks from degradation by the nuclease Mre11^74^. At least in yeast, stalled replication forks are not thought to re-localize to the nuclear periphery unless they collapse^75,76^; however, stalled forks within DNA capable of forming secondary structures can be re-localized^76^. Therefore, it is possible that, in human cells, particularly difficult substrates requiring repair, such as collapsed forks or stalled forks in difficult to replicate regions, are repositioned to facilitate this process^38^.

Another possible mechanism for overcoming stalled telomeric forks is the activation of dormant replication origins. While unperturbed telomere replication is thought to largely occur from subtelomeric regions, origins have been observed within telomeres^77,78^. Furthermore, fork stalling using APH can lead to TRF2-mediated dormant origin firing within telomeres^79^. This is a potential mechanism by which cells can ensure that telomeres are completely replicated if fork stalling occurs and cannot be restarted, which could in turn allow telomerase recruitment to the replicated telomere. However, while it is well established that replication timing of telomeres is dependent on positioning within the nucleus^80^, it is not known whether dormant origin firing would result in or require telomere re-localization.

One outstanding question from this and earlier work^38^ is the mechanism by which F-actin facilitates telomere movement. Inhibition of myosin decreased telomerase recruitment (Figure 2B, Figure S2C) and reduced movement of stalled replication forks^34^. This suggests that the motor function of myosins in actin filament movement or transport of cargo along actin fibres plays a direct role in the re-localization of stressed telomeres, although the mechanism of myosin involvement in this process remains unknown. More compelling support is the observation that telomeres on F-actin are significantly less mobile (Figures 5J-K). This implies that a yet unknown factor(s) can facilitate binding between telomeres and F-actin, which would allow for re-localization of telomeres as has been seen in this work (Figure 4C) and another study^38^. Both ARP2/3 and WASP associate with replication forks and facilitate the accumulation of RPA at stalled forks^43^, suggesting a more direct role in the DDR outside their regulation of nuclear F-actin polymerization. While the latter study did not examine telomeres specifically, it is likely that this process also occurs at stalled telomeric forks to maintain their integrity, which may consequently result in telomere tethering to F-actin. Whether ARP2/3 and WASP facilitate telomere tethering directly or indirectly remains to be elucidated.

Overall, our data reveal a mechanism by which telomerase responds to telomeric replication stress in order to maintain telomere integrity and length. We have shown that telomerase recruitment to telomeres requires ATR- and mTOR-mediated nuclear actin filament formation, and that telomerase interactions with telomeres occur in the vicinity of these filaments. We hypothesize that critically stressed telomeres are restrained by F-actin, allowing scanning telomerase molecules to find these telomeres and extend their length in lieu of DNA replication (Figure 7B). Concurrently, mildly stressed telomeres are re-localized to the nuclear periphery for resolution and complete telomere replication, which may be followed by telomerase recruitment for productive telomere extension (Figure 7C). Overall, this work suggests that there is a tight balance between replication stress and telomere maintenance, where the natural stress generated from telomere replication is exploited to facilitate telomerase recruitment, while telomerase can also protect cells from critical replication stress and sudden telomere loss.

## ACKNOWLEDGEMENTS

We thank Jens Schmidt and Tom Cech for providing hTERT- and TRF2-labelled HeLa cells used for live cell imaging, and Andrew Robinson for purification of the JF646-HaloTag ligand. We are grateful to Scott Cohen for many discussions, some of which were actually helpful. We thank the Children’s Medical Research Institute and the ACRF Telomere Analysis Centre, supported by the Australian Cancer Research Foundation, for providing access to microscopy equipment. Cell cycle analysis was performed at the Westmead Scientific Platforms, which are supported by the Westmead Research Hub, the Cancer Institute New South Wales, the National Health and Medical Research Council and the Ian Potter Foundation. This project was supported by grants from the Australian Research Council (Discovery Project grant DP190103572) and Perpetual Foundation (IPAP2022/0347). A.H. was supported by the Profield Foundation and the Neil and Norma Hill Foundation.

## AUTHOR CONTRIBUTIONS

A.H., M.K. and T.M.B. designed the project and conceived hypotheses; A.H., M.K., N.M.M., D.R.N., and K.W. performed experiments; A.H., and W.E.H. prepared custom scripts for Fiji; all authors analyzed data and interpreted results; A.H., and T.M.B. wrote the manuscript with input from all authors.

## METHODS

### EXPERIMENTAL MODEL AND STUDY PARTICIPANT DETAILS

#### Cell lines

HEK293T (T. Adams, CSIRO), A549 (American Type Culture Collection), HT1080 (American Type Culture Collection), HCT116 (G. Chenevix-Trench), HeLa-EM2-11ht (J. Schmidt, Michigan State University) and CRISPR-modified HeLa-EM2-11ht cells expressing FLAG-Halo-tagged hTERT and HA-mEOS3.2-tagged TRF2^58^ were all grown in a humidified 37 °C incubator with 5% CO_2_. HCT116 cells were cultured in McCoy’s 5A medium, supplemented with 10% fetal bovine serum (FBS) and 2 mM L-glutamine. All other cells were cultured in Dulbecco’s modified Eagle’s medium (DMEM) supplemented with 10% FBS. Cell Bank Australia validated all cell line identities by short-tandem-repeat profiling and tested all cells for mycoplasma.

## METHOD DETAILS

### Cell culture

Where necessary, cells were treated with chemical inhibitors for 16 hr prior to harvesting or sample preparation. Inhibitors are listed in the key resource table, with concentrations provided in figure captions. APH treatment (1.5 μM) was performed for 30 min immediately prior to harvesting or sample preparation, while maintaining chemical inhibition where appropriate. Dimethylsulfoxide (DMSO) was used as a vehicle control for all non-treated controls.

### siRNA transfection

For knockdown by siRNA, cells were transfected with 30 pmol siRNA and 5.5 μl Lipofectamine RNAiMAX in 200 μl Opti-MEM. Cells were grown for 48 hr after transfection to allow for knockdown. siRNAs and catalogue numbers are listed in the key resource table.

### Exogenous actin and actin chromobody transfection

Nuclear WT and R62D mutant actin expression vectors were generated from pmCherry-C1 actin-3×NLS P2A mCherry and pmCherry-C1 R62D actin-3×NLS P2A mCherry vectors (D. Mullins, University of California)^32^, respectively, by removing mCherry to enable hTR-telomere SABER FISH. BFP- and NLS-tagged actin chromobody (nuclear-actin-ChB) was generated from the NLS-GFP-actin chromobody vector (Chromotek) by replacing green fluorescent protein (GFP) with BFP. For expression of either exogenous actin or the nuclear-actin-ChB, cells were transfected with 2.5 μg plasmid DNA and 3.75 μl Lipofectamine 3000 in 250 μl Opti-MEM. Cells were grown for 48 hr after transfection to allow for expression.

### Flow cytometry

Following treatment or transfection, cells were harvested with trypsin, pelleted at 500 g for 5 min and washed twice with phosphate buffered saline (PBS). Cells were resuspended in 1 ml PBS, before adding 5 ml of cold 70% ethanol dropwise while vortexing. Cells were left at 4 °C at least overnight to fix. The day before analysis, cells were repelleted and washed twice with PBS. Cells were resuspended in fresh propidium iodide (PI) solution (50 μg/ml PI, 0.5 μg/ml RNase A, 5% Triton X-100 in PBS) and left overnight at 4 °C to stain. Cells were then analyzed using an LSRII flow cytometer and FACSDiva software (Becton Dickinson). Cell cycle profiles were generated using FlowJo analysis software (Version 10.8).

### SABER FISH for hTR and telomeres, with immunofluorescence

FISH against hTR and telomeric DNA, with or without immunofluorescence (IF), was adapted from existing hTR-telomere FISH^8,40^ and SABER (<u>s</u>ignal <u>a</u>mplification <u>b</u>y <u>e</u>xchange reaction) FISH protocols^41^. SABER probes were extended by primer extension reaction (1X PBS, 10 mM MgSO_4_, 6 mM dNTPs [dATP, dCTP, dTTP], 8 U/μl Bst LF polymerase, 2 μM target hairpin, 5 μM target probe) at 37 °C for 2 or 6 hr, for telomere or hTR probes, respectively. Reactions were heat inactivated at 80 °C for 20 min, evaluated by agarose gel electrophoresis, and then stored at −20 °C until required. All probe and hairpin sequences are listed in Table S1.

Cells were seeded on poly-L-lysine coated coverslips for 24 hr prior to transfection, inhibitor treatment or fixation. Cells were labelled with 20 μM EdU for 15-30 min immediately prior to fixation (with subsequent APH treatment where appropriate). Cells were washed once with PBS before permeabilization with 0.5% IGEPAL (in PBS) for 10 min, then fixed with 2% paraformaldehyde (PFA, in PBS) for 15 min. Cells were washed twice with MilliQ water, before further permeabilization with 1:1 methanol:acetone solution for 10 min. Cells were washed once with PBS, then EdU was labelled with AF488 using the Click-iT® Plus reaction kit. Cells were then washed once with PBS before gradient ethanol dehydration (70%, 90% and 100% ethanol) for 2 min each and brief air drying. Probe mastermix was prepared by diluting extended hTR and telomere probes 1:100 in RNAse-free water, before denaturing at 90 °C for 2 min, followed by cooling on ice for 3 min. For each coverslip, 1 μl probe mastermix was added to 30 μl FISH hybridization buffer (2X SSC [0.3 M NaCl, 30 μM Tri-sodium citrate, pH 7], 10% dextran sulphate, 2 mM vanadyl-ribonucleoside complex, 0.02% bovine serum albumin (BSA), 1 μg/μl *E. coli* tRNA, 50% deionized formamide) before pipetting onto slides and covering with coverslips containing cells. Slides were then heated at 80 °C for 5 min before hybridizing overnight at 37 °C in a humidified chamber. Coverslips were then washed twice with FISH wash 1 (2× SSC, 50% formamide, 0.1% sodium dodecyl sulfate [SDS]) for 30 min, three times with FISH wash 2 (4× SSC, 0.1% Tween-20) for 5 min, and once with PBS briefly before allowing to air dry. Coverslips were placed onto slides containing imager probes (1 μM in PBS) and incubated in a humidified chamber at 37 °C for 1-1.5 hr. Coverslips were washed with pre-warmed (37 °C) PBS for 15 min, twice for 5 min, and then rinsed twice with MilliQ water before air drying. Coverslips were mounted with Prolong Gold with DAPI and left overnight to cure.

For IF-FISH, the same protocol was followed until the EdU labelling step; EdU was instead labelled with AF350. Following this, the coverslips were blocked with ABDIL buffer (20 mM Tris [pH 7.5], 2% BSA, 0.2% fish skin gelatin, 120 mM NaCl, 0.1% Triton X-100, 0.1% sodium azide) in a humidified chamber at RT for 1 hr. Anti-PCNA antibody was then diluted in ABDIL (1:1000) and added to coverslips, which were incubated overnight in a humidified chamber at 4 °C. Coverslips were washed with PBS for 5 min three times, before incubating with AF488-labelled secondary antibody diluted in ABDIL (1:2000) for 1 hr at RT in a humidified chamber. Coverslips were washed three times with PBS for 5 min, re-fixed with 2% PFA for 15 min, washed twice with MilliQ water, before continuing with gradient ethanol dehydration and the remaining steps outlined above. Finally, coverslips were mounted with Prolong Gold.

Fixed samples were imaged using an AxioImager Z.2 microscope (Carl Zeiss), using a ×63/1.4NA oil-immersion objective, appropriate filters, an Axiocam 506 monochromatic camera (Carl Zeiss) and Zen Blue Pro v.2.3.69.01015 (Carl Zeiss). Samples were imaged with 13 Z-stacks at 0.25 μm intervals using consistent exposure times between all treatments and experiments. Pixel intensity histograms were adjusted equally across figure panels for presentation only.

### Actin chromobody imaging and FISH/IF-FISH

Cells were seeded on poly-L-lysine coated coverslips for 24 hr prior to transfection with nuclear-actin-ChB. Samples were left for 48 hr, before treating cells with 1.5 μM APH for 1 hr to promote F-actin formation. Cells were then fixed with ice-cold 4% PFA for 15 min, washed with PBS, permeabilized with 0.5% Triton X-100 (in PBS) for 10 min and washed with PBS before ethanol dehydrating. Cells were then prepared for FISH or IF-FISH as above, excluding EdU steps, and mounted with Prolong Gold before being left to cure overnight. Samples were imaged by super-resolution microscopy (described below).

### Phalloidin staining

Cells were seeded on poly-L-lysine coated coverslips for 24 hr prior to staining. Cells were washed once with PBS, then fixed with ice-cold 4% PFA for 15 min, washed with PBS, permeabilized with 0.5% Triton X-100 (in PBS) for 10 min and washed with PBS. Cells were stained using phalloidin (fluorescein isothiocyanate labelled), diluted to 2% in PBS, overnight at 4 °C in a humidified chamber. Coverslips were then washed four times with PBS for 5 min each, before gradient ethanol dehydration (70%, 90% and 100% for 2 min each), air drying, and then mounting with Prolong Gold with DAPI. Samples were imaged by super-resolution microscopy (described below).

### Preparation of nuclear and cytoplasmic extracts

Cells were washed with PBS, harvested with trypsin, pelleted at 500 *g* for 5 min and washed once with PBS. 1 x 10^6^ cells were aliquoted into a fresh microcentrifuge tube for each sample, pelleted and then resuspended in ∼20 volumes (200 μl) of hypotonic buffer (10 mM HEPES-KOH [pH 8], 10 mM KCl, 1 mM MgCl_2_, 1 mM dithiothreitol [DTT]). Cells were placed on ice for 5-10 min to swell, as observed by microscopy, before adding 1/20 volume of 10% Triton X-100 to lyse cell membranes. Tubes were mixed by inversion and kept on ice to observe cell membrane lysis by microscopy. Nuclei were then pelleted at 1000 *g* for 1 min at 4 °C. The cytoplasmic fraction was collected in a separate tube, and the nuclei were washed twice with 1 ml hypotonic buffer. Fractionation purity was determined by western blot using vinculin and histone H2A as cytoplasmic and nuclear markers, respectively. 4× cell equivalents of nuclei were loaded relative to cytoplasmic fractions for all blots.

### Western blot

Cells were harvested with trypsin and washed with PBS. Cell lysates were prepared at 1 × 10^4^ cells/μl in 4× LDS (106 mM Tris-HCl, 141 mM Tris-base, 2% lithium dodecyl sulphate, 10% glycerol, 0.22 mM Brilliant Blue G250, 0.175 mM Phenol Red) supplemented with 50 mM DTT and 2% Benzonase for 15 min at RT. Lysates were denatured at 68 °C for 10 min, before loading 7 × 10^4^ cell equivalents onto Nu-PAGE 4-12% gradient Bis-Tris gels and electrophoresed in MES SDS running buffer at 150 V for 70 min. Proteins were transferred to Amersham 0.45 μm nitrocellulose (GE Healthcare) in transfer buffer (25 mM Tris-Base, 192 mM glycine, 10% methanol) at 30 V overnight at 4 °C. Membranes were blocked in 5% skim milk powder or BSA in TBST (15 mM Tris-HCl, 4.6 mM Tris-Base, 137 mM NaCl, 0.1% Tween-20) for 1 hr before incubation with gentle rotation overnight at 4 °C with primary antibodies diluted in either 5% skim milk powder or BSA in TBST. Membranes were washed three times for 10 min with TBST, before probing for 1 hr at RT with secondary antibodies diluted in 5% skim milk powder in TBST. Membranes were washed three times for 10 min with TBST, before developing signal using either Amersham ECL or ECL prime detection reagent. Blots were imaged using a BioRad ChemiDoc™ MP imaging system.

### Live cell imaging

HeLa cells expressing FLAG-Halo-tagged hTERT and HA-mEOS3.2-tagged TRF2^58^ were seeded on sterile 35 mm FluoroDish plates (Coherent Scientific) with cover glass bottoms. Cells were allowed to settle overnight before transfecting with nuclear-actin-ChB. Cells were left for 8 hr and then synchronized by addition of 2 mM thymidine for 15 hr, releasing for 9 hr, and addition of 2 mM thymidine for 16 hr. Cells were released into S phase with fresh DMEM (+ 10% FBS) for 3-5 hr before labelling FLAG-HaloTag-hTERT. HaloTag ligand was prepared by coupling with Janelia Fluor 646^81^ following the ligand manufacturer’s instructions, followed by purification using liquid chromatography/mass spectrometry (LC/MS) on a C18 column (3.5 µm, 21.2 mm x150 mm) with a 3-90% acetonitrile/water gradient at 2 mL/min and detection at 656 nm. JF-646-Halo ligand (50 nM) in DMEM was added to cells for 1-5 min, before washing the cells three times with DMEM and returning cells to the incubator in DMEM (+ 10% FBS) for 15 min. Cells were washed 3 times with DMEM, before addition of DMEM without phenol red (+ 10% FBS) prior to imaging. Cells were imaged by super-resolution microscopy (described below).

### Super-resolution microscopy

Super-resolution imaging was performed using an LSM 880 AxioObserver confocal laser scanning fluorescent microscope (Carl Zeiss) fitted with a super-resolution Airyscan detector using a Plan-Apochromat ×63/1.4NA M27 oil-immersion objective and ZEN Black 2.3 Pro v14.0.20.201 (Carl Zeiss). All samples/cells were imaged using ‘Resolution versus sensitivity’ mode, appropriate filter sets, unidirectional scanning, 1×1 binning and frame averaging (4 frames for fixed samples, 2 frames for live cell imaging).

Fixed samples were imaged using 405 nm (0.2% excitation power, 790 detector gain), 488 nm (1.16% excitation power, 790 detector gain), 561 nm (0.5% excitation power, 913 detector gain) and 633 nm (2.66% excitation power, 860 detector gain) lasers. Live cells were imaged at 37 °C, 20% O_2_ and 5% CO_2_, using 405 nm (0.2% excitation power, 790 detector gain), 561 nm (0.5% excitation power, 913 detector gain) and 633 nm (5% excitation power, 903 detector gain) lasers. Cells were briefly (10-20 sec) exposed to the 405 nm laser immediately before imaging to convert HA-mEOS3.2-TRF2 from green to red wavelength. Imaging frame time was ∼0.5 sec per channel, with all channels imaged consecutively for 250-1000 frames.

## QUANTIFICATION AND STATISTICAL ANALYSIS

### Automated FISH/IF-FISH image analysis

Microscopy images were converted into extended projections of Z-stacks using ZEN blue desk software v2.3 (Carl Zeiss) and exported as TIFF files for analysis using CellProfiler v2.2.0^82^. Individual nuclei and foci were identified using intensity-based thresholding strategies. Individual foci were considered overlapping with another focus if at least 25% of the first focus was contained within the second. Triple co-localization events (hTR, telomere and PCNA) were counted if 25% of an hTR focus was within a telomere focus, and if either also overlapped (any amount) with a PCNA focus. At least 100 S phase cells were quantified per treatment, with each treatment performed in triplicate.

### Measurement of minimum distance to nuclear periphery or F-actin

Measurement of the minimum distance to the nuclear periphery or F-actin was performed using custom scripts for Fiji v1.53q^83^. Individual nuclei, foci, and co-localizations were identified using CellProfiler v2.2.0 as above, and all identified objects were exported as binary images for import into Fiji. F-actin was detected using the FilamentDetector plugin (v2.0.0) for Fiji. For each focus/co-localization, the distance between the centroid and each point along the nuclear periphery or the nuclear F-actin network was calculated, with the lowest value returned.

### Classification of live cell imaging recruitment events

Prior to analysis, live cell imaging data was processed using a custom Fiji script to improve foci clarity. CZI files were imported into Fiji, and each channel was separated for frame averaging and processing. Images were averaged on a 10-frame basis, before background subtraction (2 pixel rolling ball radius by sliding paraboloid) and gaussian blur (sigma 1). Channels were merged and exported as TIFF files for analysis in Imaris software v9.9.1. Recruitment events were classified manually where hTERT and TRF2 foci overlapped at least 25% for a minimum of 2 consecutive averaged frames (∼30 sec). Recruitment events were classified as ‘observed’ only if one of the two foci was observed in at least the two previous consecutive averaged frames prior to recruitment; all other events were classified as ‘previously recruited’. Recruitment events and individual hTERT or TRF2 foci were sub-classified based on the distance to a visible actin fibre. Foci and events were classified as ‘on fibre’ if one (or both) focus was at least ∼10% overlapping with F-actin. Anything under ∼10%, with no observable gap between focus and F-actin was classified as ‘adjacent’. Foci/co-localizations within ∼1.5 μm were classified as ‘proximal’, with everything else classified as ‘distal’.

### Telomere MSD analysis

Telomere tracking was performed using live cell imaging data without any processing. Data were imported into Imaris v9.9.1, where the nuclei were identified using the ‘Surfaces’ function and the nuclear-actin-ChB channel. To correct for cell shifting or migration, we utilized previously written code for MATLAB R2023b to ‘register’ and align nuclei consistently within the frame^34^. The registration operation from MATLAB was returned to Imaris, which was completed using the ‘Align image without interpolation’ function. Following registration, TRF2 foci were identified using the ‘Spots’ function in Imaris (0.5 μm focus diameter, >20 quality). Foci were then tracked using ‘Autoregressive Motion’ tracking. Tracks were generated based on ∼0.5 μm maximal distance movement between frames, 0 gaps within each track, and a minimum track length of 30 sec. The MSD was calculated for each timepoint (except T0) within each track and classified based on the intensity of nuclear-actin-ChB signal within the focal mask. All timepoints from cells within an experiment were pooled together to generate a single MSD value.

### Statistical analysis

Statistical analyses were performed using GraphPad Prism v9.4.1. One-way ANOVA with Tukey’s multiple comparison test was used for all experiments unless otherwise noted.

## KEY RESOURCES TABLE

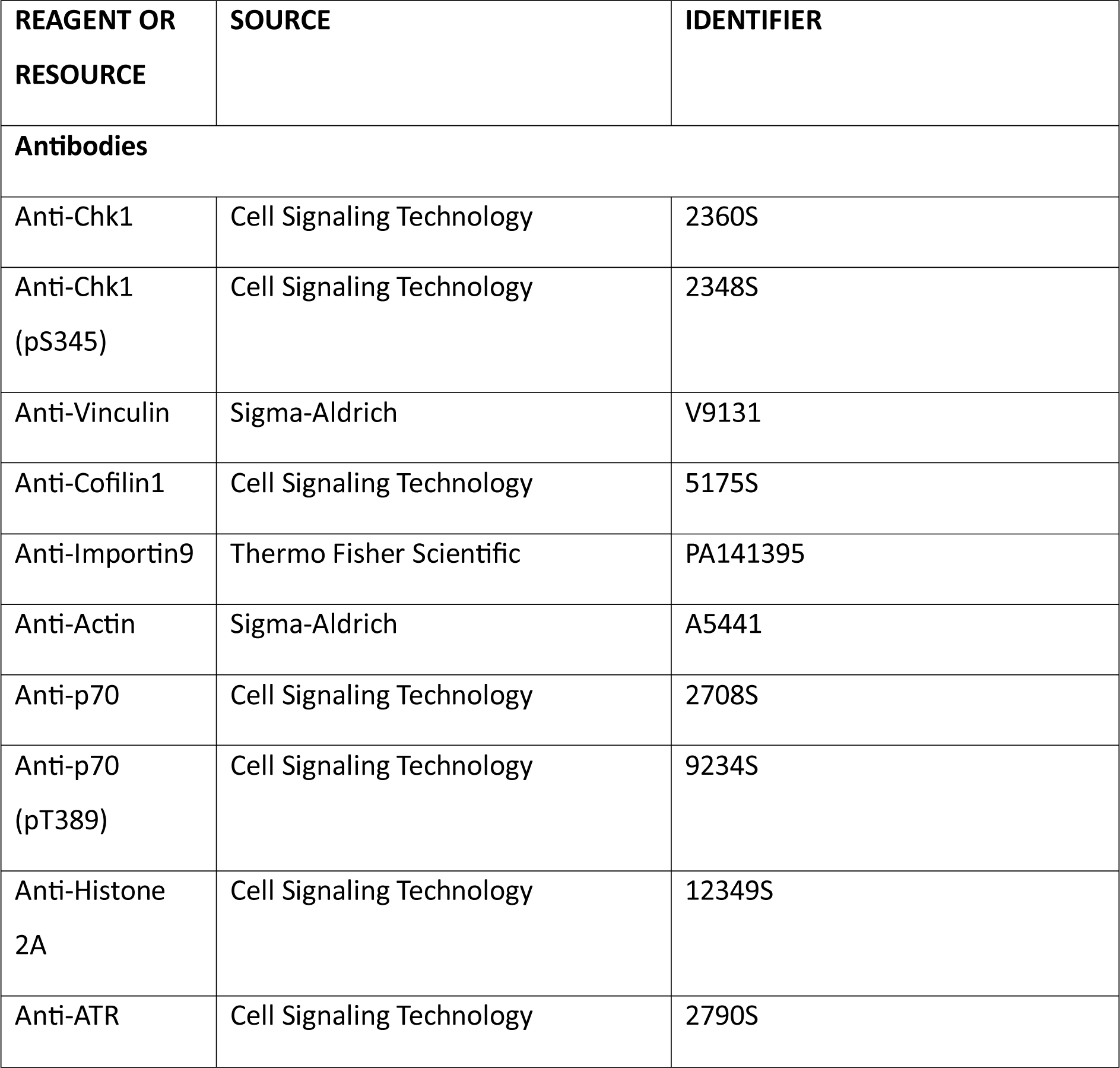
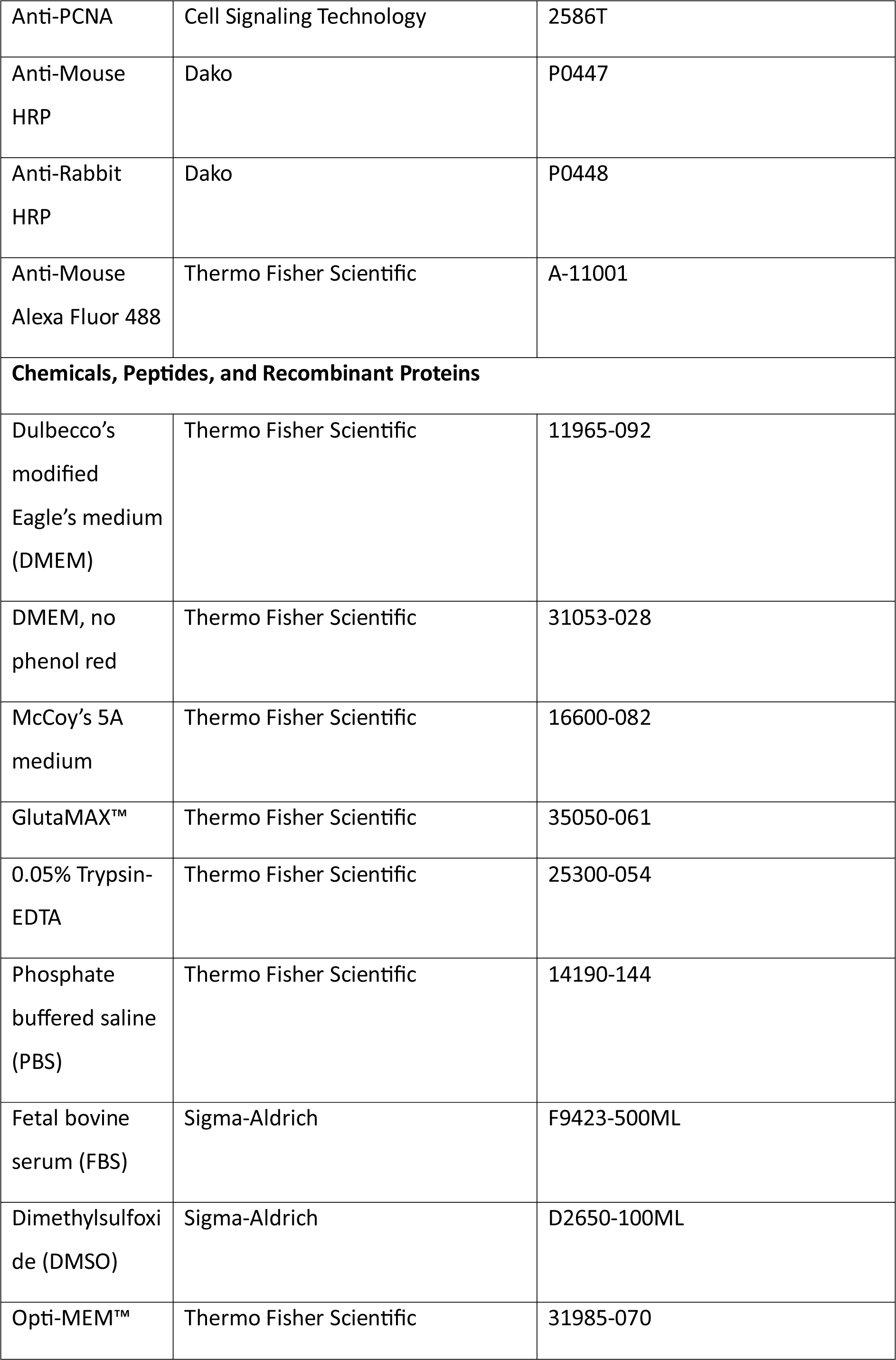
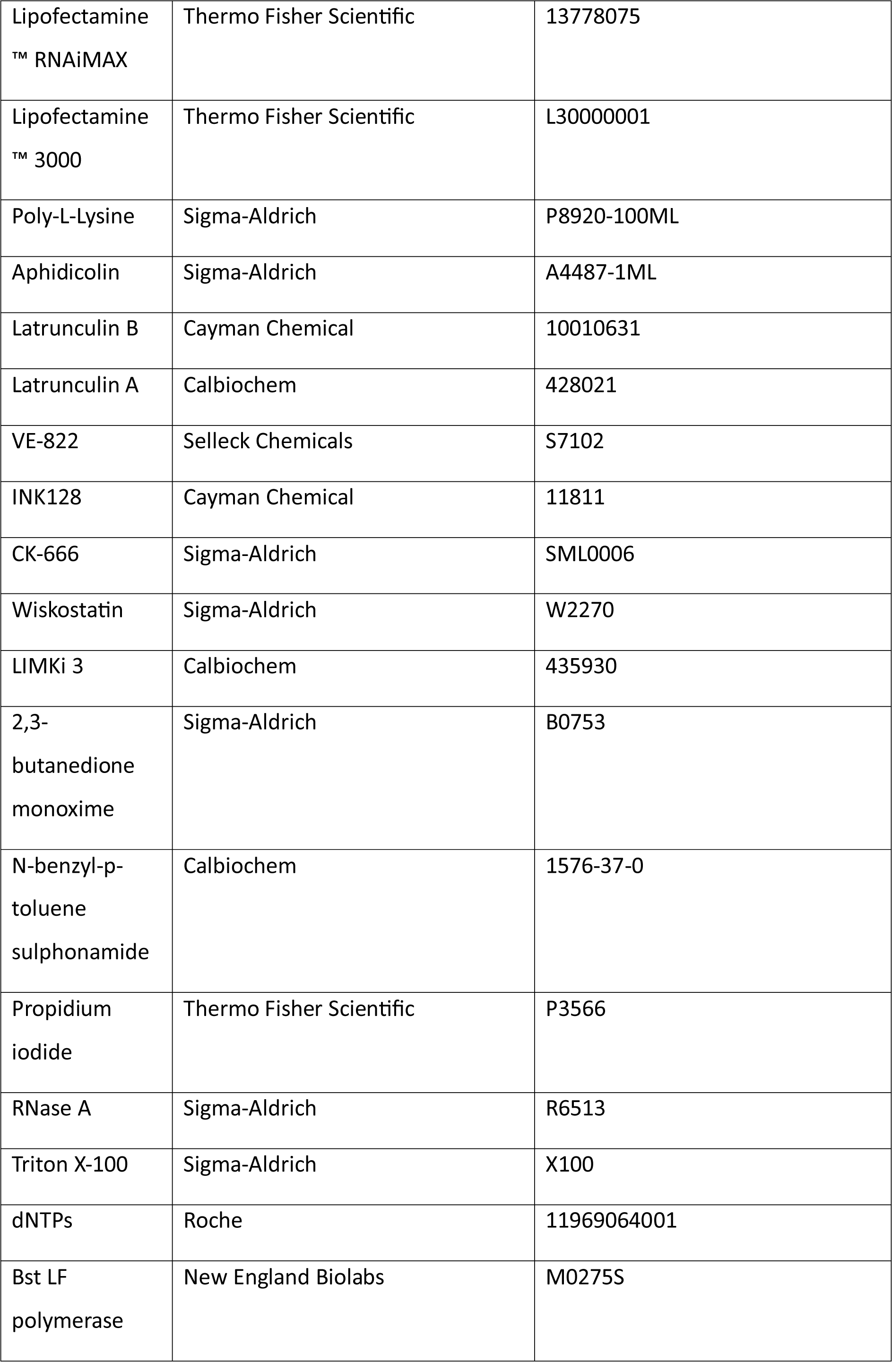
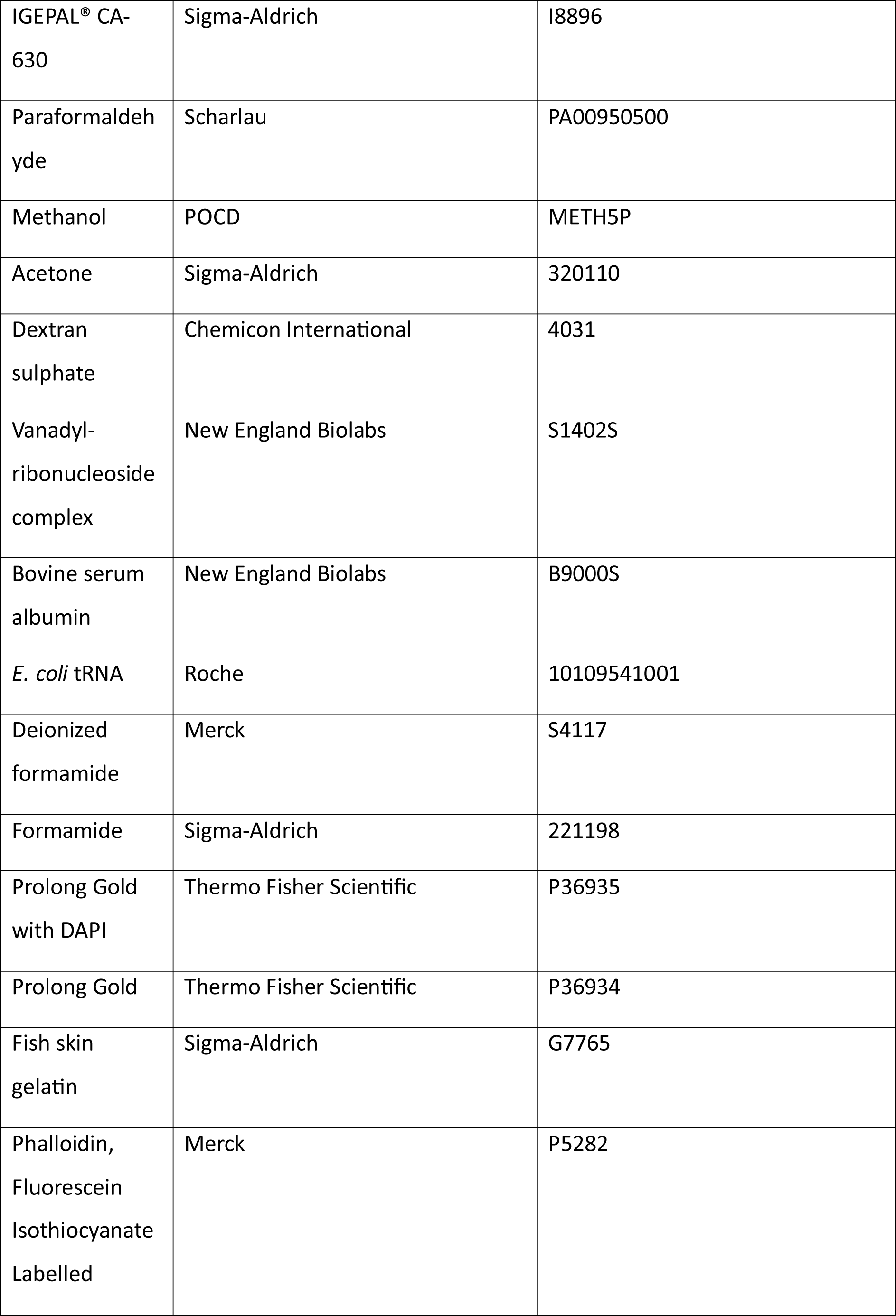
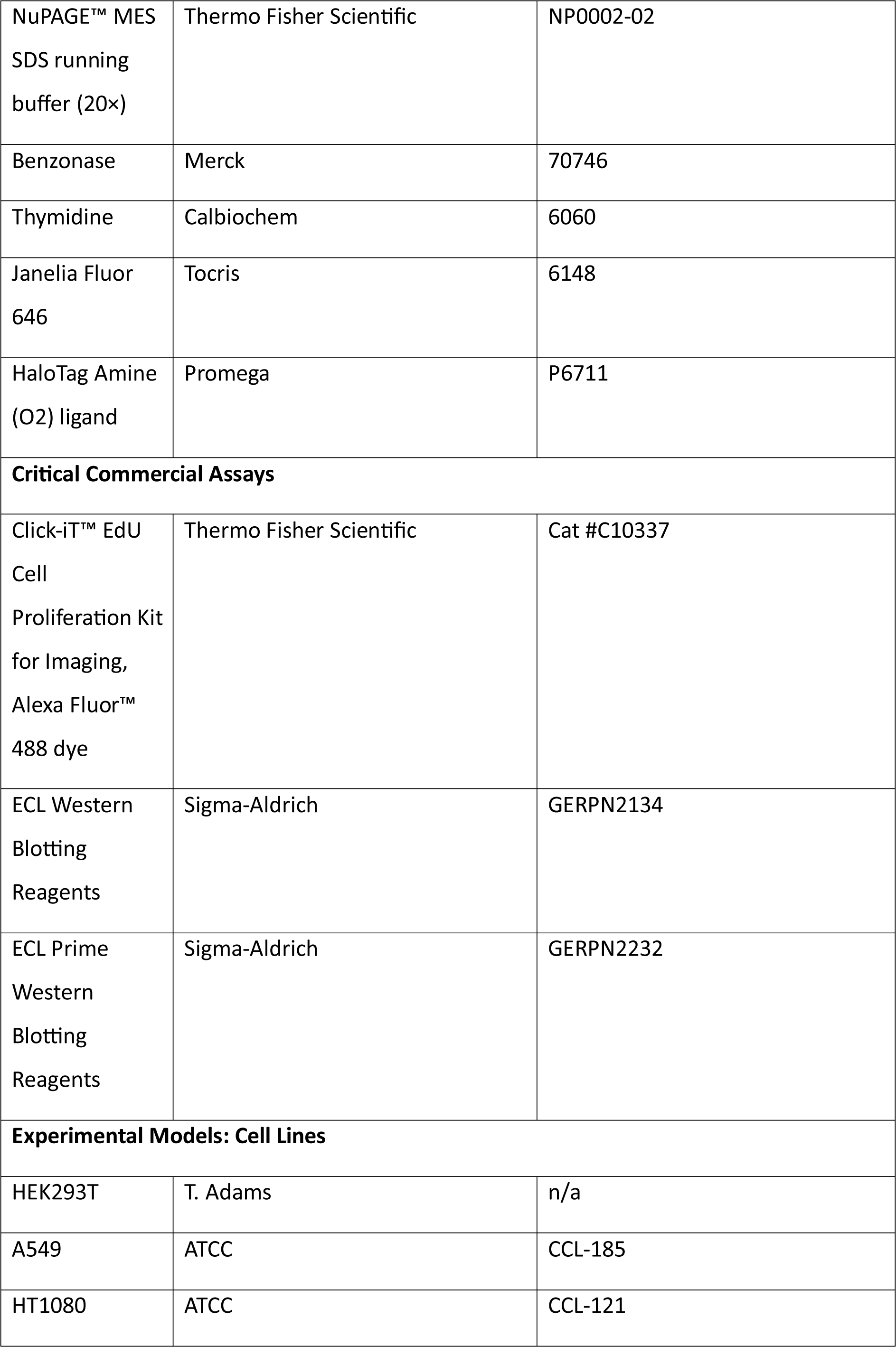
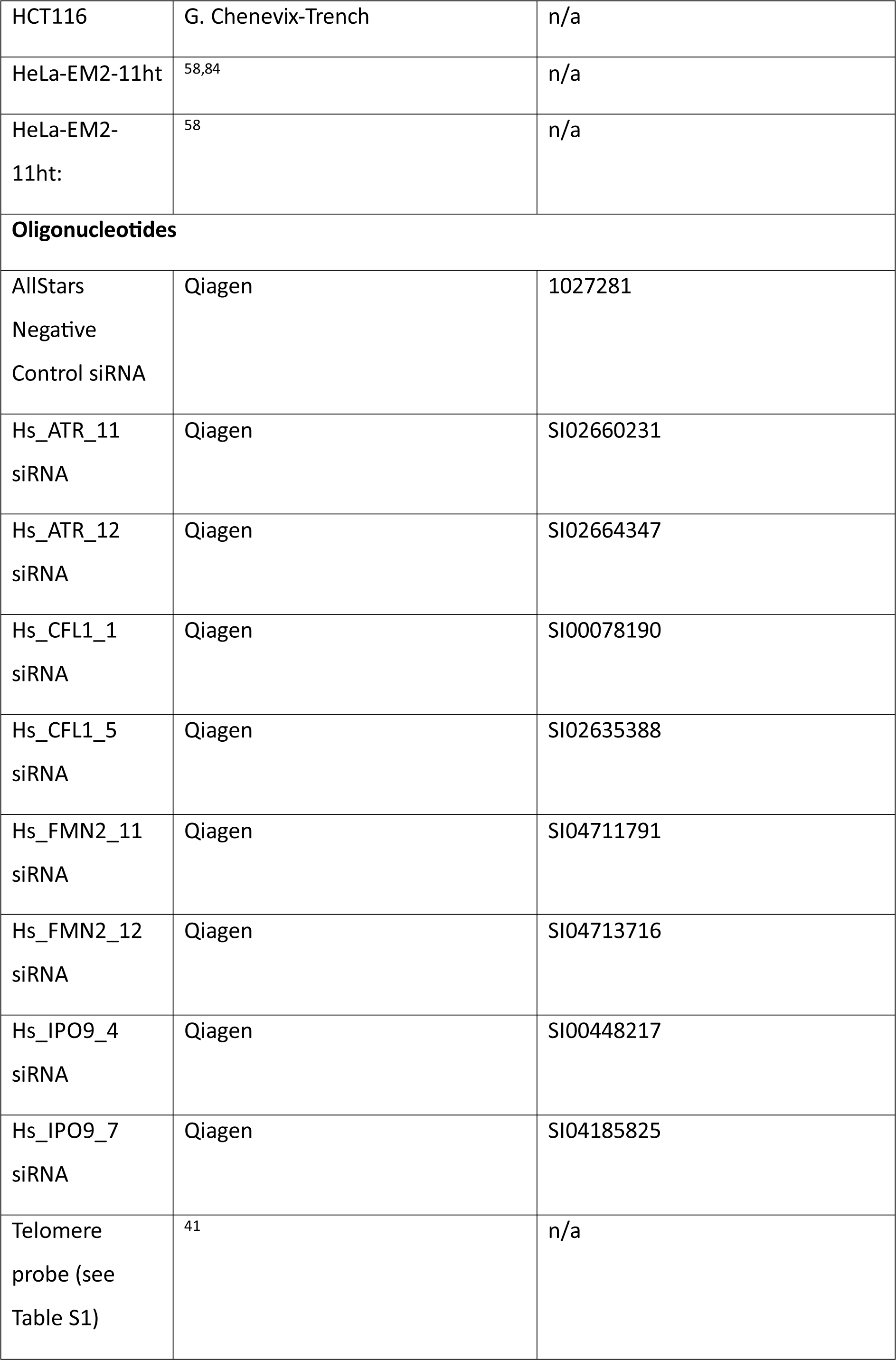
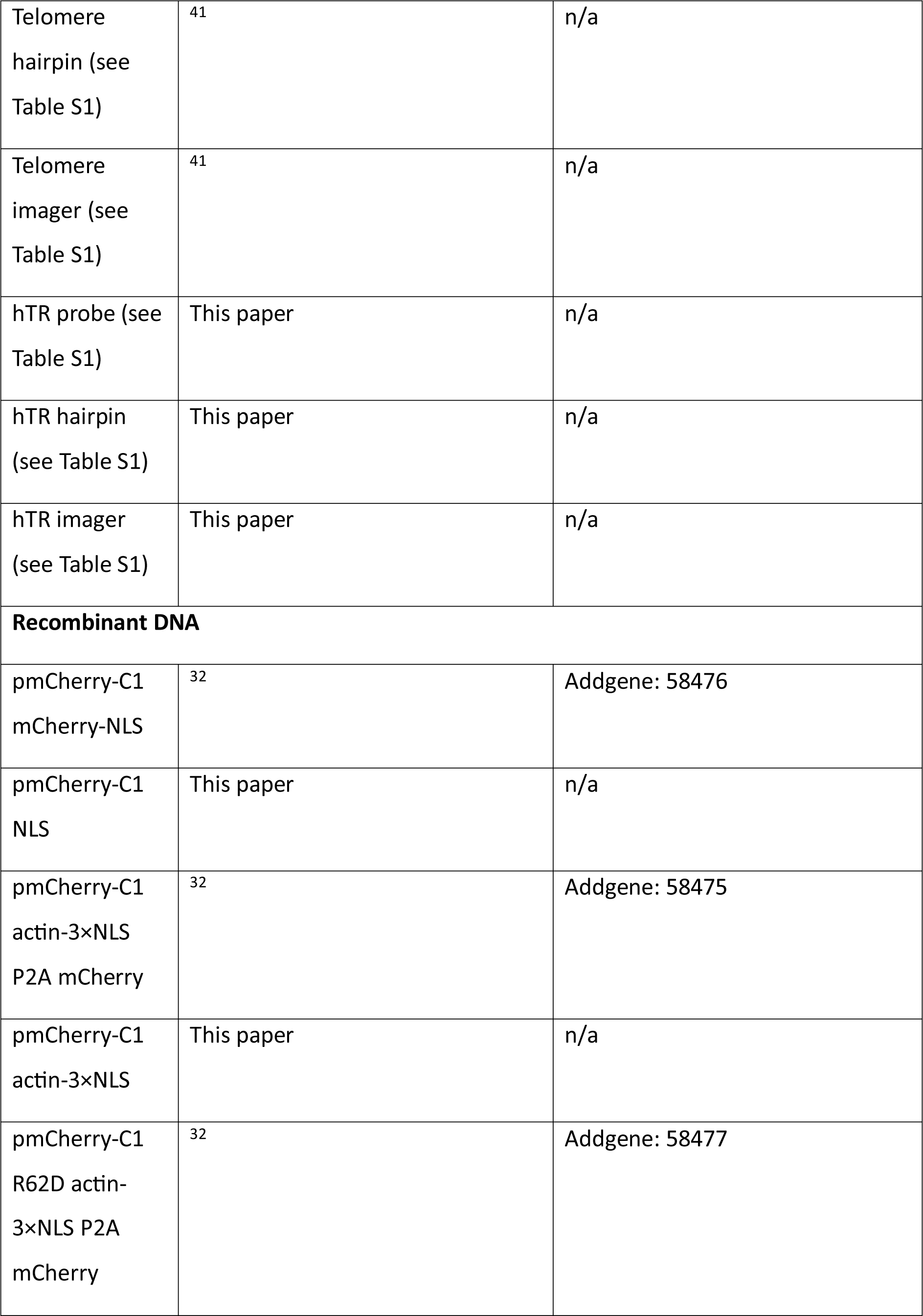
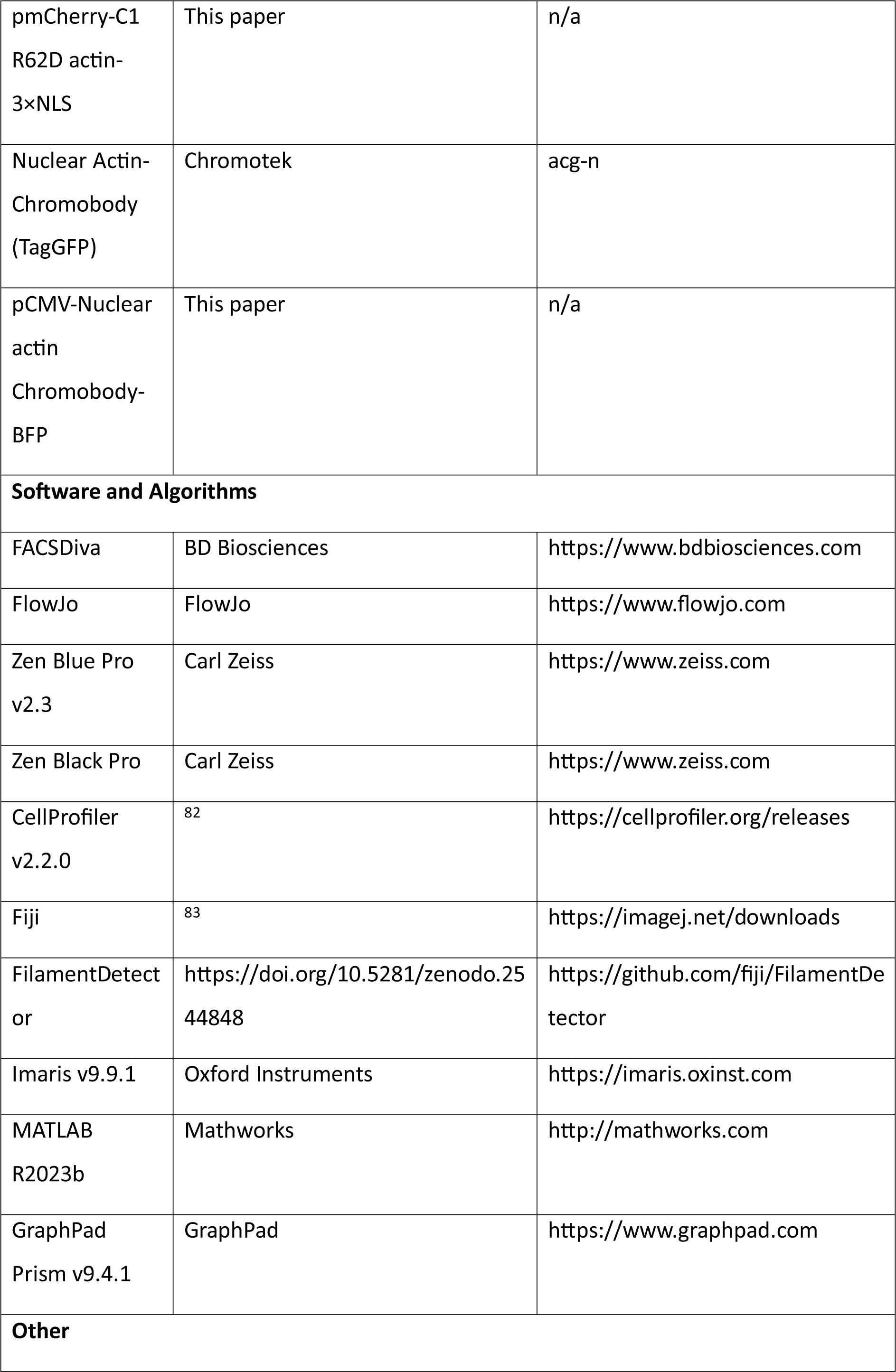
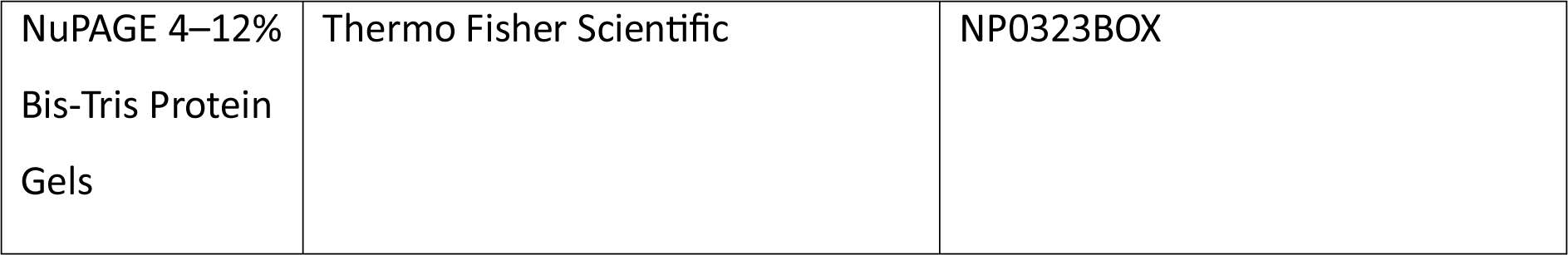

