## Supplementary data for "Nuclear actin and DNA replication stress regulate the recruitment of human telomerase to telomeres"

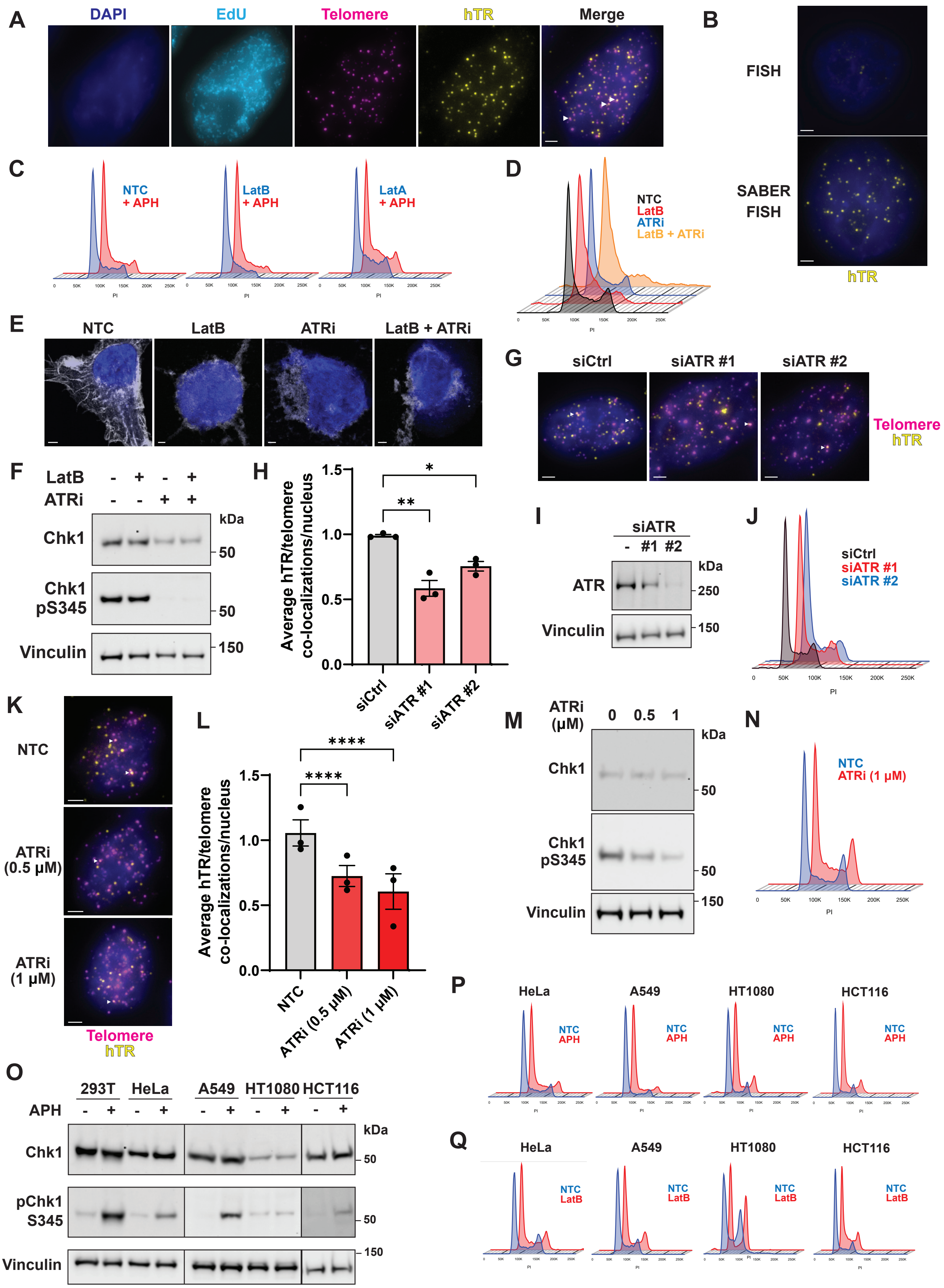
A
DAPI
EdU
Telomere
hTR
Merge
B
FISH
SABER FISH
hTR
C
NTC + APH
LatB + APH
LatA + APH
PI
D
NTC
LatB
ATRi
LatB + ATRi
PI
E
NTC
LatB
ATRi
LatB + ATRi
F
LatB - + - +
ATRi - - + +
kDa
Chk1
50
Chk1 pS345
50
Vinculin
150
H
Average hTR/telomere co-localizations/nucleus
1.5
1.0
0.5
0.0
*
**
siCtrl
siATR #1
siATR #2
G
siCtrl
siATR #1
siATR #2
Telomere
hTR
I
siATR
- #1 #2
kDa
ATR
250
Vinculin
150
J
siCtrl
siATR #1
siATR #2
PI
K
NTC
ATRi (0.5 μM)
ATRi (1 μM)
Telomere
hTR
L
Average hTR/telomere co-localizations/nucleus
1.5
1.0
0.5
0.0
****
****
NTC
ATRi (0.5 μM)
ATRi (1 μM)
M
ATRi (μM)
0 0.5 1
kDa
Chk1
50
Chk1 pS345
50
Vinculin
150
N
NTC
ATRi (1 μM)
PI
O
293T HeLa A549 HT1080 HCT116
APH - + - + - + - + - +
kDa
Chk1
50
pChk1 S345
50
Vinculin
150
P
HeLa
A549
HT1080
HCT116
NTC
APH
Q
HeLa
A549
HT1080
HCT116
NTC
LatB

**Figure S1. Inhibition of F-actin polymerization inhibits telomerase recruitment, but not DNA damage response or cell cycle progression, related to Figure 1**

(A) Example fluorescent microscopy images of modified SABER FISH protocol using probes against telomeres (pink) and hTR (yellow). All cells are stained with DAPI (blue), and S phase cells are identified with EdU (cyan) labelled using AF488. Co-localizations are indicated by white arrows in the merge panel. Scale bar, 2 µm.

(B) Comparison between standard FISH and modified SABER FISH protocols probing for hTR (yellow). Cells were stained with DAPI (blue) to visualize nuclei and imaged using identical exposure settings. Scale bar, 2 µm.

(C) Cell cycle analysis by flow cytometry of propidium iodide (PI) stained 293T cells treated with DMSO (NTC), 0.2 µM LatB or 0.05 µM LatA, ± 1.5 µM APH.

(D) Cell cycle analysis of PI-stained 293T cells treated with DMSO (NTC), 0.2 µM LatB, 0.25 µM VE-822 (ATRi), or both LatB and ATRi.

(E) 293T cells stained with phalloidin (white) and DAPI (blue) following treatment with 0.2 µM LatB, 0.25 µM ATR inhibitor VE-822 (ATRi) or LatB and ATRi together. Scale bar, 2 µm.

(F) Western blots of 293T cells treated with LatB, VE-822 (ATRi) or both LatB and ATRi, probed for Chk1 and Chk1 pS345, with vinculin as a loading control.

(G) Representative SABER FISH images of HeLa cells treated with control or ATR siRNA, stained with DAPI (blue) and probed for hTR (yellow) and telomeres (pink). Co-localizations are indicated by white arrows. Scale bar, 2 µm.

(H) Average hTR/telomere co-localizations in HeLa cells treated with control siRNA or two different ATR siRNAs.

(I) Western blot of HeLa cells treated with control or ATR siRNAs, probed for ATR and vinculin as loading control.

(J) Cell cycle analysis of PI-stained HeLa cells treated with control or ATR siRNAs.

(K) Representative SABER FISH images of HeLa cells treated with DMSO (NTC) or 0.5-1 µM VE-822 (ATRi), stained with DAPI (blue) and probed for hTR (yellow) and telomeres (pink). Co-localizations are indicated by white arrows. Scale bar, 2 µm.

(L) Average hTR/telomere co-localizations in HeLa cells treated with DMSO (NTC) or VE-822 (ATRi).

(M) Western blots of HeLa cells treated with DMSO or VE-822 (ATRi), probed for Chk1, Chk1 pS345 and vinculin as a control.

(N) Cell cycle analysis of PI-stained HeLa cells treated with DMSO (NTC) or 1 µM VE-822 (ATRi).

(O) Western blots of 293T, HeLa, A549, HT1080 and HCT116 cells treated with DMSO or 1.5 µM APH, probed for Chk1, Chk1 pS345, and vinculin as a loading control.

(P) Cell cycle analysis of PI-stained HeLa, A549, HT1080 and HCT116 cells treated with DMSO (NTC) or 1.5 µM APH.

(Q) Cell cycle analysis of PI-stained HeLa, A549, HT1080 and HCT116 cells treated with DMSO (NTC) or 0.2 µM LatB.

All bar graphs displayed as mean ± SEM; $n = 3$; ****$p<0.0001$, **$p<0.01$,*$p<0.05$.

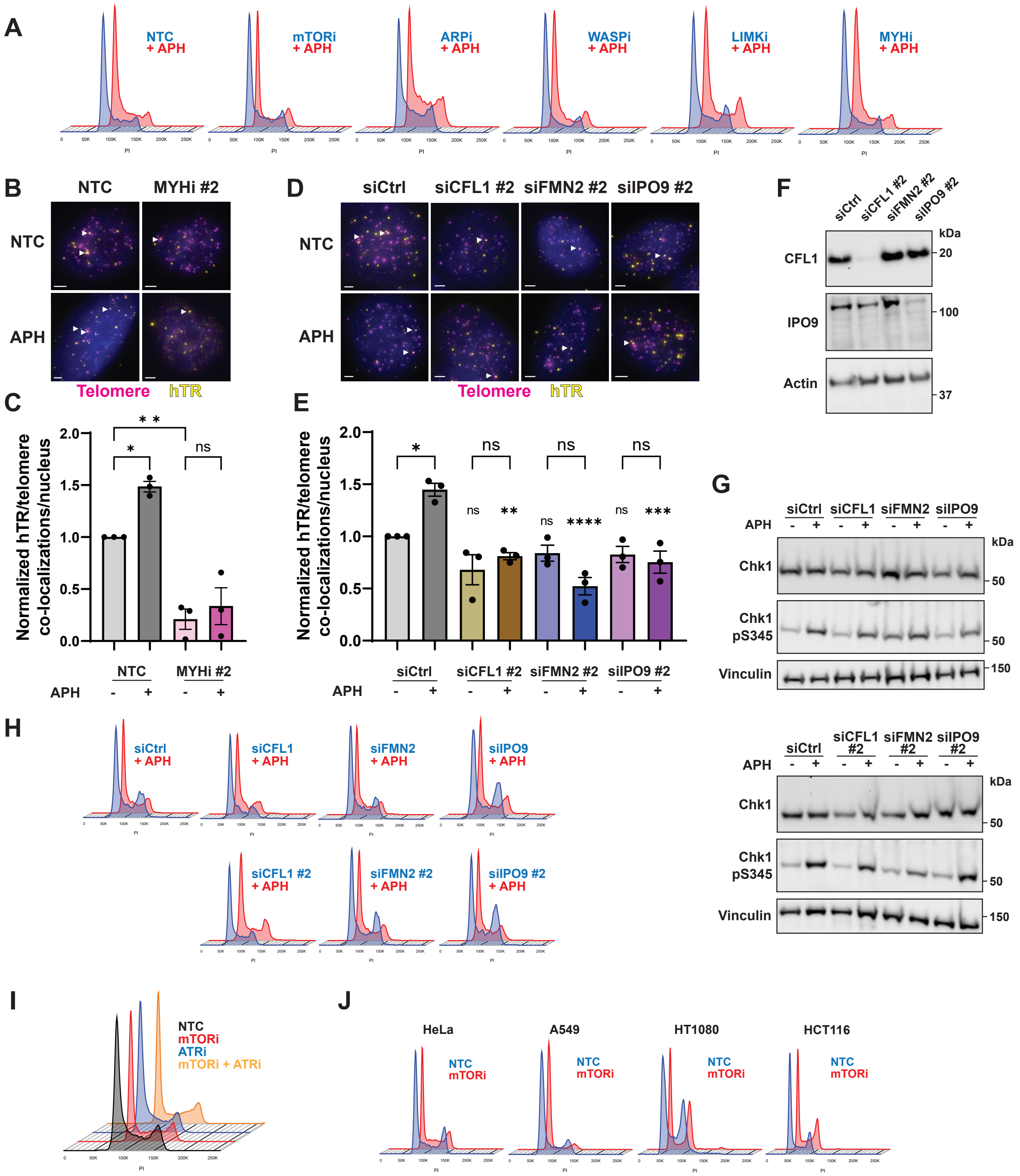
A
NTC + APH
mTORi + APH
ARPi + APH
WASPi + APH
LIMKi + APH
MYHi + APH
B
NTC
MYHi #2
NTC
APH
Telomere
hTR
C
Normalized hTR/telomere co-localizations/nucleus
* *
*
ns
NTC
MYHi #2
APH
D
siCtrl
siCFL1 #2
siFMN2 #2
siIPO9 #2
NTC
APH
Telomere
hTR
E
Normalized hTR/telomere co-localizations/nucleus
*
ns
ns
ns
ns
**
ns
****
ns
***
siCtrl
siCFL1 #2
siFMN2 #2
siIPO9 #2
APH
F
siCtrl
siCFL1 #2
siFMN2 #2
siIPO9 #2
kDa
CFL1
20
IPO9
100
Actin
37
G
siCtrl
siCFL1
siFMN2
siIPO9
APH
kDa
Chk1
50
Chk1 pS345
50
Vinculin
150
siCtrl
siCFL1 #2
siFMN2 #2
siIPO9 #2
APH
kDa
Chk1
50
Chk1 pS345
50
Vinculin
150
H
siCtrl + APH
siCFL1 + APH
siFMN2 + APH
siIPO9 + APH
siCFL1 #2 + APH
siFMN2 #2 + APH
siIPO9 #2 + APH
I
NTC
mTORi
ATRi
mTORi + ATRi
J
HeLa
A549
HT1080
HCT116
NTC
mTORi

**Figure S2. Known nuclear F-actin regulators are important for telomerase recruitment to telomeres, not the DNA damage response or cell cycle progression, related to Figure 2**

(A) Cell cycle analysis by flow cytometry of PI-stained 293T cells treated with DMSO (NTC), 0.2 µM INK128 (mTORi), 200 µM CK-666 (ARPi), 5 µM wiskostatin (WASPi), 10 µM LIMKi 3 (LIMKi) or 10 mM BDM (MYHi), ± 1.5 µM APH.

(B) Representative images of 293T treated with DMSO (NTC) or 10 nM BTS (MYHi #2) ± 1.5 µM APH, probed for hTR (yellow) and telomeres (pink) with DAPI staining (blue). Co-localizations are indicated by white arrows. Scale bar, 2 µm.

(C) Normalized hTR/telomere co-localizations in 293T cells treated with DMSO (NTC) or BTS (MYHi #2) ± APH.

(D) Representative SABER FISH images of 293T cells probed for hTR (yellow) and telomeres (pink) with DAPI (blue) staining, following transfection with control siRNA, or a second siRNA against cofilin 1 (CFL1), formin 2 (FMN2) or importin 9 (IPO9), ± 1.5 µM APH treatment. Co-localizations are indicated by white arrows. bar, 2 µm.

(E) Normalized telomerase presence at telomeres in 293T cells transfected with control siRNA, or a second siRNA against CFL1, FMN2 or IPO9, ± APH. Significance for each siRNA treated sample is expressed relative to the appropriate control (siCtrl ± APH). siCtrl data are the same as presented in Figure 2E.

(F) Western blot of 293T cells treated with control siRNA, or a second siRNA against CFL1, FMN2 or IPO9, probing for CFL1, IPO9 and actin as a control. Note: FMN2 western blot not shown due to lack of specific antibody.

(G) Western blot of 293T cells transfected with control siRNA, or two siRNAs against each of CFL1, FMN2 or IPO9, ± APH, probing for Chk1, Chk1 pS345, and vinculin as a loading control.

(H) Cell cycle analysis of PI-stained 293T cells transfected with control, CFL1, FMN2 or IPO9 siRNA, ± APH.

(I) Cell cycle analysis of PI-stained 293T cells treated with DMSO (NTC), INK128 (mTORi), VE-822 (ATRi) or mTORi and ATRi in combination.

(J) Cell cycle analysis by flow cytometry of HeLa, A549, HT1080 and HCT116 cells treated with DMSO (NTC) or INK128 (mTORi).

All bar graphs displayed as mean ± SEM; n = 3; ****$p<0.001$, ***$p<0.001$,**$p<0.01$,*$p<0.05$.

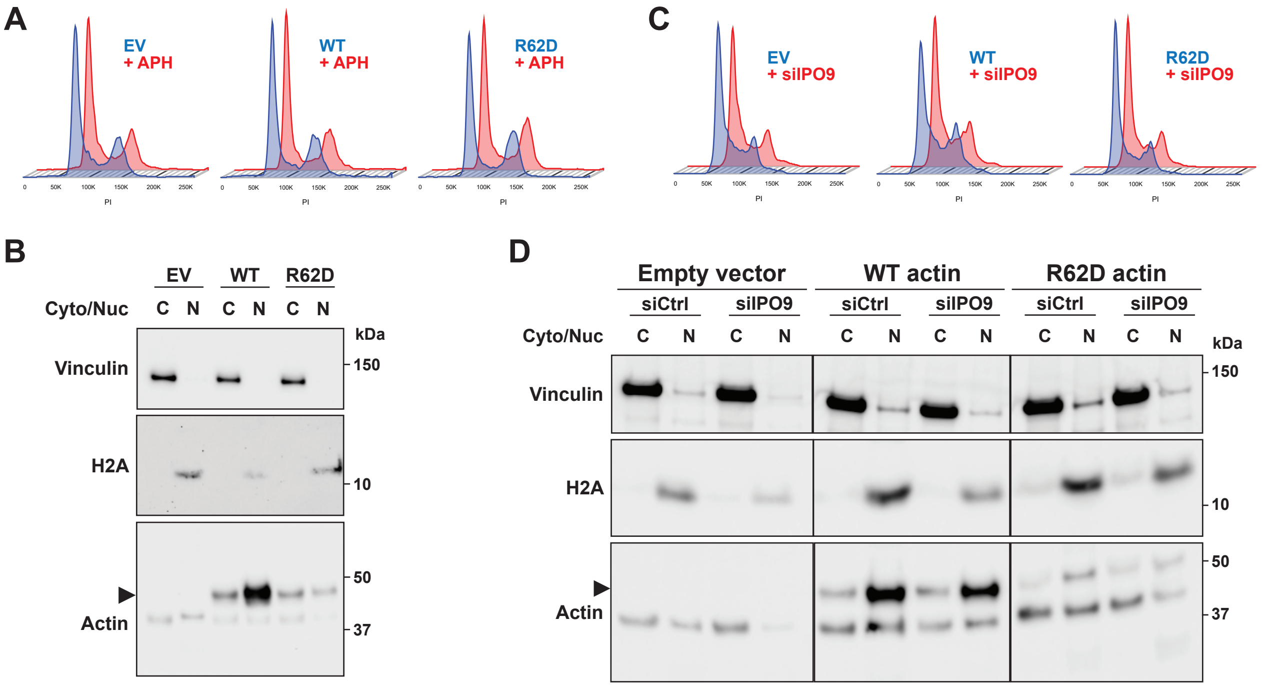


**Figure S3. Validation of nuclear WT and mutant (R62D) actin expression, related to Figure 3**

(A) Cell cycle analysis by flow cytometry of PI-stained 293T cells transfected with empty vector (EV) or vectors encoding WT or mutant (R62D) actin, ± APH treatment.

(B) Western blot analysis of 293T cells transfected with empty vector (EV) or vectors encoding WT or mutant (R62D) actin. Nuclei were separated from cytoplasmic content via cell fractionation prior to analysis. Blots were probed with vinculin (cytoplasmic marker), H2A (nuclear marker) and actin. Exogenous actin is tagged with a 3×NLS, and is indicated by a black arrow. 4× cell equivalents of nuclei were loaded relative to cytoplasmic fractions.

(C) Cell cycle analysis of PI-stained 293T cells transfected with vectors encoding WT or mutant (R62D) actin, with or without IPO9 siRNA. Empty vector (EV) and control siRNA were used as controls.

(D) Western blot analysis of 293T cells transfected with vectors encoding WT or mutant (R62D) actin, with or without IPO9 siRNA. Empty vector (EV) and control siRNA were used as controls. Nuclei were separated from cytoplasmic content via cell fractionation prior to analysis. Blots were probed with vinculin (cytoplasmic marker), H2A (nuclear marker) and actin. Exogenous actin is tagged with a 3×NLS, and is indicated by a black arrow. 4× cell equivalents of nuclei were loaded relative to cytoplasmic fractions.

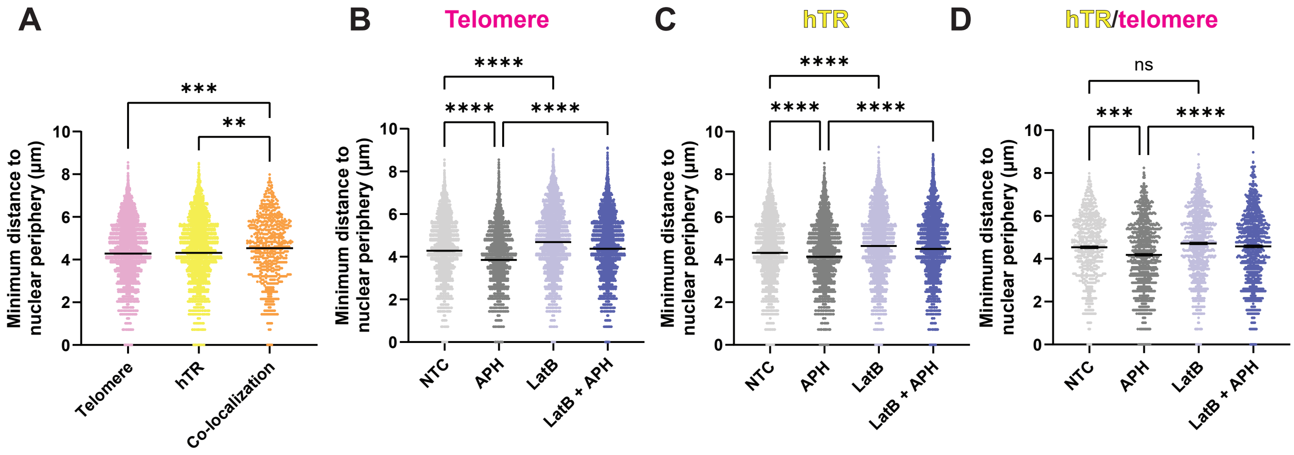


**Figure S4. Movement of telomeres and telomerase toward nuclear periphery following replication stress, related to Figure 4**

(A) Data from Figure 4B displayed as individual values representing the minimum distance of telomeres (pink), hTR (yellow), or hTR/telomere co-localizations (orange) from the nuclear periphery in 293T cells. n = 14756 (telomeres), 6123 (hTR) and 624 (co-localizations).

(B-D) Data from Figures 4C-E displayed as individual values representing the minimum distance of telomeres (B), hTR (C), or hTR/telomere co-localizations (D) from the nuclear periphery in 293T cells treated ± 1.5 µM APH and 0.2 µM LatB. Telomeres: n = 14756 (NTC), 19360 (APH), 11030 (LatB) and 12708 (LatB + APH). hTR: n = 6123 (NTC), 8981 (APH), 9839 (LatB) and 7207 (LatB + APH). Co-localizations: n = 624 (NTC), 910 (APH), 988 (LatB) and 864 (LatB + APH).

All lines on graphs represent mean ± SEM; ****p<0.0001, ***p<0.001,**p<0.01.

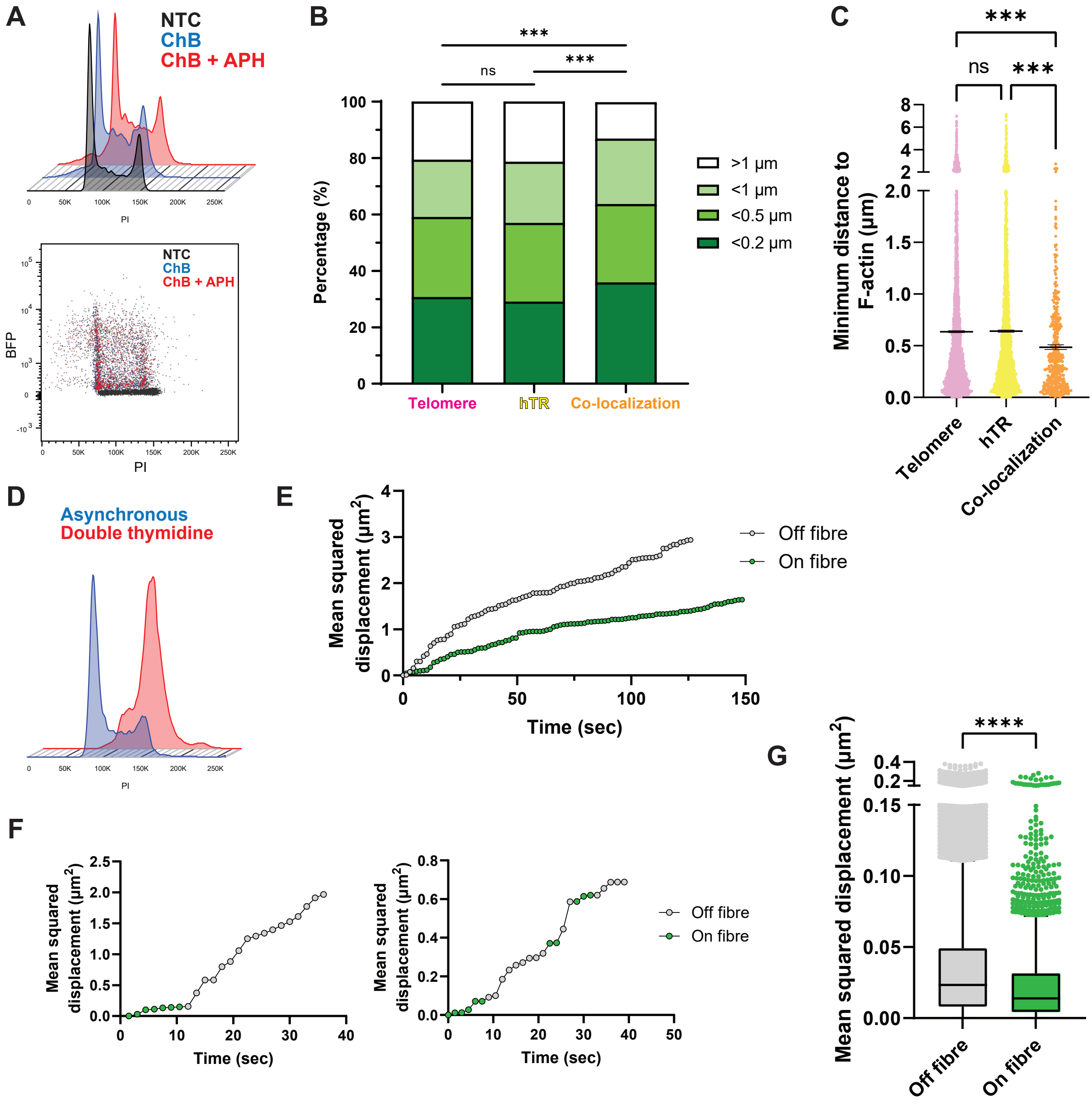
A
NTC
ChB
ChB + APH
PI
BFP
B
Percentage (%)
ns
***
***
Telomere
hTR
Co-localization
>1 µm
<1 µm
<0.5 µm
<0.2 µm
C
Minimum distance to F-actin (µm)
ns
***
***
Telomere
hTR
Co-localization
D
Asynchronous
Double thymidine
PI
E
Mean squared displacement (µm$^2$)
Time (sec)
Off fibre
On fibre
F
Mean squared displacement (µm$^2$)
Time (sec)
Off fibre
On fibre
G
Mean squared displacement (µm$^2$)
****
Off fibre
On fibre

**Figure S5. Recruitment of telomerase to telomeres in proximity to nuclear F-actin, related to Figure 5**

(A) Top: Cell cycle analysis of PI-stained HeLa cells transfected with the nuclear-actin-ChB (ChB) ± APH treatment. Bottom: BFP intensity in PI-stained HeLa cells. Untransfected (NTC) cells were used as a control.

(B) Pooled data from Figure 5C displayed as percentages based on distance from nuclear F-actin. Statistics were performed using $\chi^2$ tests. n = 7384 (telomeres), 5929 (hTR) and 434 (co-localizations).

(C) Pooled data from Figure 5C displayed as individual data points with mean ± SEM. n = 7384 (telomeres), 5929 (hTR) and 434 (co-localizations).

(D) Cell cycle analysis of PI-stained HeLa cells. Cells were synchronized by double thymidine block, followed by release into S phase for 5 hours. Asynchronous cells were used as a control.

(E) Further examples of the cumulative mean squared displacement (MSD) of telomere (TRF2) foci in relation to F-actin. MSD is plotted over time for a focus which is not on F-actin (grey) or one that remained associated with F-actin (green).

(F) Examples of cumulative mean squared displacement (MSD) of two telomere (TRF2) foci that moved off and on F-actin fibres over the course of imaging. MSD is plotted over time for foci which transition between being off (grey) or on F-actin (green).

(G) Pooled data from Figure 5K plotted as individual MSD values of telomere (TRF2) foci from live cell imaging experiments. MSD was calculated for each telomere on a frame-by-frame basis, and each MSD value was classified based on the intensity of nuclear-actin-ChB signal within the focus region. n = 24505 (Off fibre) and 3647 (On fibre). Data is displayed as a Tukey boxplot.

****p<0.0001, ***p<0.001.

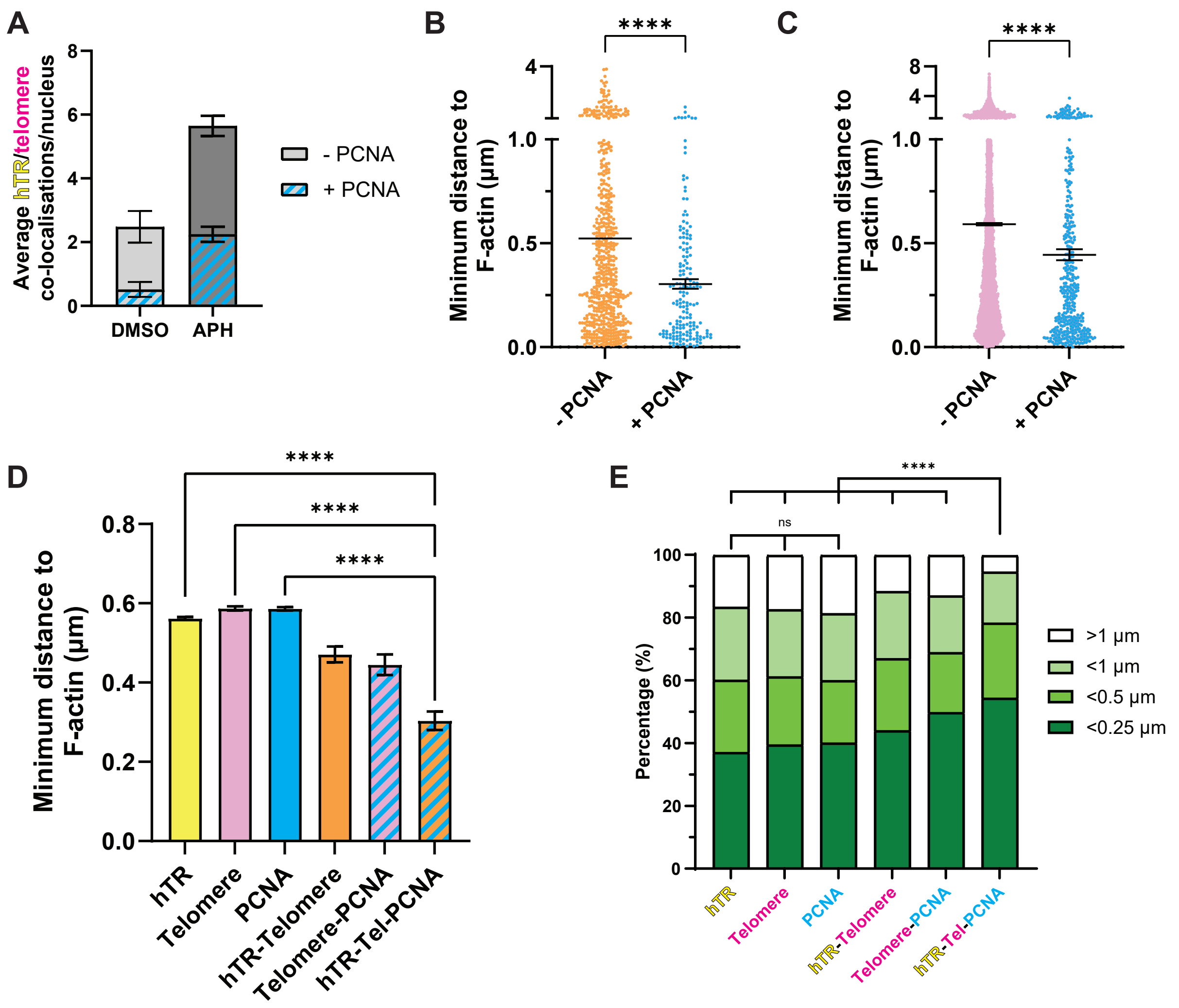


**Figure S6. Telomerase recruitment occurs at F-actin with stalled replication forks, related to Figure 6**

(A) Data from Figure 6E shown as average hTR/telomere co-localizations which also co-localized with PCNA foci in 293T cells treated ± APH. Data displayed as mean ± SEM; n = 3.

(B-C) Pooled data from Figure 6H displayed as individual data points with mean ± SEM. Data are pooled from >200 nuclei; n = 562 (hTR/telomere), 171 (hTR/telomere/PCNA), 15098 (telomeres) and 418 (telomere/PCNA).

(D) Pooled data from Figure 6H, including individual hTR and PCNA foci. Data are not separated based on co-localizations (i.e. hTR data includes all hTR foci, including those which overlap with telomeres and/or PCNA, etc). Data are pooled from >200 nuclei; n = 25421 (hTR), 15579 (telomere), 23129 (PCNA), 726 (hTR/telomere), 418 (telomere/PCNA) and 171 (hTR/telomere/PCNA).

(E) Pooled data from Figure S6D displayed as percentages based on distance from nuclear F-actin. Statistics were performed using $\chi^2$ tests.

****p<0.0001,*p<0.05.

**Table S1.** Related to STAR Methods. SABER FISH probes and oligonucleotides used for primer extension reactions.

| **Name** | **Sequence (5' – 3')** **PER repeats in red, hairpin sequences underlined** | **Repeat no. (Kishi *et al.,* 2019)** | **Purification** |
|---|---|---|---|
| Telomere Probe | TAACCCTAACCCTAACCC TAACCCTAACCCTAACCC TAACCCTAACCCTTACATCATCATACATCATCAT | 27 | Standard desalting |
| Telomere Hairpin | ACATCATCATGGGCCTTTTGGCCCATGATGATGTATGATGATG /3InvdT/ | 27 | HPLC |
| Telomere Imager | /5Cy3/ ATGATGATGTATGATGATGT | 27 | HPLC |
| hTR Probe | CTCCGTTCCTCTTCCTGCGG CCTGAAAGGCCTGAACCTCG CCCTCGCCCCCGAGAGTTACAACTTAACACAACTTAAC | 28 | Standard desalting |
| hTR Hairpin | ACAACTTAACGGGCCTTTTGGCCCGTTAAGTTGTGTTAAGTTG /3InvdT/ | 28 | HPLC |
| hTR Imager | /5Cy5/ GTTAAGTTGTGTTAAGTTGT | 28 | HPLC |

**Supplementary Videos 1+2**

Examples of telomerase recruitment to telomeres in HeLa cells. Representative video of a CRISPR-modified HeLa cell expressing HA-mEOS3.2-tagged TRF2 (pink) and FLAG-HaloTag-hTERT (visualized using JF646-HaloTag ligand; yellow). Cells were transfected with nuclear-actin-ChB to visualize nuclear actin (green) and synchronized by double thymidine block before 3-5 hr release into S phase and imaging via super-resolution Airyscan microscopy. Raw images were averaged on a 10-frame basis before background subtraction and guassian blur. Time is shown as minutes:seconds relative to the first frame of the video.

**Supplementary Video 3**

Nuclear F-actin impacts telomere mobility under replication stress. Representative video of a CRISPR-modified HeLa cell expressing HA-mEOS3.2-tagged TRF2 (pink) and transfected with nuclear-actin-ChB to visualize nuclear actin (green). Cells were synchronized by double thymidine block before 3-5 hr release into S phase and imaging via super-resolution Airyscan microscopy. Blue and white boxes indicate telomeres on or off F-actin, respectively. Time is shown as minutes:seconds relative to the first frame of the video.
